# Dynamic winter microbial communities shape nitrogen cycling potential in Arctic tundra soils

**DOI:** 10.64898/2026.04.07.717072

**Authors:** Stephanie Turner, Dominik Merges, Emil Alexander Sherman Andersen, Niki Ines Walter Leblans, Ellen Dorrepaal, Sara Hallin, Karina Engelbrecht Clemmensen

## Abstract

Arctic winters are long and cold and have traditionally been considered a period of limited biological activity. However, the seasonal dynamics of microbial community composition and functional potential during winter remain poorly understood. Here, we investigated taxonomic (bacteria, fungi, archaea) and functional (fungal guilds and nitrogen cycling genes) dynamics throughout a full year at two Arctic tundra heath sites with contrasting snow regimes. A steep drop in microbial abundances in early to mid-winter, likely linked to freeze-thaw events, coincided with shifts in soil pH and elevated community turnover. Saprotrophic and root-associated fungi were more abundant in the cold-season, while inorganic nitrogen cycling groups were more abundant in summer and declined toward winter despite high bacterial abundance. This indicates sustained organic matter cycling during the winter and expanded inorganic nitrogen cycling in the summer. Functional gene ratios further suggested a higher early-winter nitrogen loss potential via nitrous oxide and greater late-winter nitrogen retention. Site-specific differences in snow regime altered the timing and magnitude of these dynamics. Together, our results demonstrate that winter represents a critical and dynamic period for microbial community restructuring with important implications for nitrogen turnover in Arctic tundra soils.

## Introduction

Arctic winters are long and cold and have traditionally been viewed as a period of biological dormancy, during which most organisms are inactive or experience high mortality. In soils, freezing causes the formation of ice crystals, limited liquid water availability, and increased osmotic pressure, which can damage microbial cells and lead to cell death and lysis (Mazur, 1984; Öquist et al., 2009). However, this paradigm is challenged by studies demonstrating that microorganisms can survive, grow and remain metabolically active at subzero temperatures (Drotz et al., 2010; Elberling & Brandt, 2003; Natali et al., 2019). Because snow acts as a thermal insulator, snow cover and soil temperatures are interconnected. Increasing snow depth buffers soils from air temperature extremes, allowing mild soil temperatures and potentially enabling soil microbial communities to survive and remain active. Although cold season activities are often low, the cold season (i.e. non-growing season) makes up a substantial part of the year for Arctic ecosystems, and cumulative processes can contribute substantially to the annual biogeochemical budgets and ecosystem-scale processes including greenhouse gas emissions (Gao et al., 2022; Natali et al., 2019). Despite these advances, very little is known about dynamics of microbial community composition and functional potential during winter, and how seasonal restructuring of microbial communities may have implications for elemental cycling.

Among the few studies available, microbial abundance patterns are inconsistent, partly depending on the methodological approach, whereas microbial community composition show seasonal changes (Buckeridge et al., 2013; Schostag et al., 2015). Microbial community turnover appears to be highest during the transition periods with freeze-thaw cycles in fall and in spring, whereas winter communities seem to be relatively stable (Poppeliers et al., 2022). Seasonal dynamics of Arctic microbial communities might also be linked to the seasonal variation in nitrogen (N) availability. Arctic soils are generally N-poor as they receive low inputs of N via atmospheric deposition (Dentener et al., 2006). Further, microorganisms compete for N both within and among bacteria, fungi, and archaea, and with plants. Plants display a seasonality in N uptake (McKane et al., 2002), thereby also affecting the year-round dynamics of N. In spring, a transition from high concentrations of dissolved organic N before snow melt, followed by a pulse of ammonium after thaw, and a pulse of nitrate shortly after, was reported for an alpine and an Arctic tundra (Broadbent et al., 2021; McLaren et al., 2018). This suggests a succession of different microbial functional groups, i.e. an increase of nitrifiers, converting ammonia to nitrate, after thaw, potentially followed by an increase in denitrifiers and nitrate ammonifiers using nitrate as a substrate. During the growing season, microorganisms in Arctic tundra soils have shown to be N limited (Sistla et al., 2012). In a tundra heath, gross nitrification rates were found to be stable during summer, although gross N mineralization and N availability dropped, likely due to plant uptake and N leaching, indicating an increasing N limitation (Koranda & Michelsen, 2024). Schostag et al. (2015) identified up to 24% of the active community to be N_2_ fixing bacteria in a mesic meadow tundra, which likely counteracted limitation during summer. In fall, when plants senesce and add fresh litter to the soil system, saprotrophic organisms thrive, increasing organic matter (OM) degradation and potentially N mineralization, causing another pulse in ammonium and inorganic N cycling communities (McLaren et al., 2018). In contrast to the growing season, there is very little information about microbial communities involved in N turnover in tundra soils during the long cold season. The few available studies suggest that dissolved organic N availability peaks during mid-winter (Broadbent et al., 2021; Buckeridge et al., 2013; Schostag et al., 2015), which could be a result of ongoing OM mineralization in snow-covered soils, while plant N uptake is more limited. While the high availability of N might stimulate microbial inorganic N cycling processes, cold temperatures likely suppress microbial growth, which could create a situation of increased risks of N losses. These responses, however, also depend on whether N uptake by plants and their closely associated mycorrhizal fungi would continue during the cold season. Recent evidence from Arctic tundra heath indeed suggested sustained plant N uptake potential throughout the Arctic winter (Andersen et al., in revision), which may further imply that perennial Arctic plants maintain their support of root-associated mycorrhizal fungi throughout the winter. Thus, to understand N dynamics through the Arctic winter, it is necessary to analyze a broad suite of microbial functional groups in the soil at high temporal resolution.

Here, we followed bacterial, fungal and archaeal communities in the soils of two sub-Arctic tundra sites with contrasting snow regime by monthly sampling throughout a full year, with the aim to evaluate dynamics of abundance, community composition, and functional capacity of the three microbial groups, as well as their synchronies, during the Arctic winter. While several studies have focused on the effect of snow regimes on microbial communities and activities during the subsequent spring and summer (Broadbent et al., 2021; Buckeridge & Grogan, 2010; Kolstad et al., 2021; Ricketts et al., 2016; Semenova et al., 2016), information is scarce on how the snow regime influences microbial community composition and functional capacities throughout the winter itself. We further aimed to evaluate the potential implications of microbial community dynamics for the turnover of N. We hypothesized that (1) bacterial and archaeal abundances decline in fall and increase in spring, while fungal abundances remain stable year-round due to dominance of perennial mycelia, and (2) microbial community turnover is highest in spring and fall linked to drastic changes in environmental conditions. Lastly, we hypothesized that (3) these patterns are also reflected in functional groups: i) Root-associated fungal guilds are expected to remain stable throughout the year, while saprotrophic litter decomposing fungi peak in fall, and molds and yeasts peak in spring, coupled to the turnover of microbial biomass and the release of inorganic N, and ii) bacterial and archaeal N functional gene abundances, reflecting the genetic potential for N-cycling, are expected to decrease in fall, whereas the release of inorganic N in spring first stimulates nitrifiers, followed by denitrifiers and nitrate ammonifiers, and, as N becomes increasingly limited during summer, diazotrophs that fix N_2_ will increase.

## Material and Methods

### Sampling sites

Over the course of one year from July 2019 until August 2020, soil samples were taken at two sub-Arctic mixed tundra heath sites close to Abisko (68°21’N, 18°49’E) and Vassijaure (68°25’N, 18°15’E) in Northern Sweden. The two sites differ in microclimate, i.e., Abisko is dryer, with a mean annual precipitation of 347 mm, compared to 859 mm at Vassijaure (SMHI, 1991-2020). The mean annual temperatures at Abisko and Vassijaure were 0.3°C and −0.5°C (SMHI, 1991-2020), respectively. The vegetation at both sites was characterized by a mix of evergreen and deciduous shrubs and graminoids, and the dominant vascular plant species included *Empetrum hermaphroditum* Hagerup, *Betula nana* L., *Vaccinium uliginosum* L., *Andromeda polifolia* L., and *Carex* spp. The vegetation cover in Vassijaure had a higher proportion of mosses (77 ± 2 %, mean ± standard error) compared to Abisko (63 ± 3 %) (Fig. S1).

Sampling procedure, site characteristics, and experimental setup are described in detail in Andersen et al. (in revision). Briefly, sampling took place approximately every four weeks resulting in a total of 15 time points. Each site had five plots (‘blocks’) that were at least 7 m apart, and each plot included 15 patches, one for each sampling time point, at least 1 m apart to minimize lateral disturbance between sampling occasions. At each time point, a soil core of approximately 11 cm was taken from one patch per block and used for the labelling experiment described in Andersen et al. (in revision). After removing this soil core, two smaller soil cores (2.2 cm diameter, ca. 1-2 cm long) were taken horizontally from within the core hole at opposite sides at approximately 5-7 cm soil depth and transported to the lab on dry ice. These two smaller cores were collected for the microbial analyses of the present study and were stored at −80°C until further processing.

### Environmental and soil data

Data acquisition of environmental parameters is described in detail in Andersen et al. (in revision). In brief, air temperatures were obtained from meteorological stations (Abisko and Katterjåkk, SMHI), located 1 km (Abisko) and 3 km (Vassijaure) away from the sites. Soil temperatures were measured via loggers (EM50 data loggers, Decagon Devices Ltd., WA, USA) at 5 cm depth at each block. Snow cover was measured before each sampling at the respective soil patch. Soil moisture and chemistry parameters were measured on part of the larger soil core at each harvest. Soil moisture was determined gravimetrically. Soil pH was measured in a soil-MilliQ water-suspension (1:5) with a pH electrode (InLab® Expert NTC 30, Mettler Toledo, OH, USA). All soil N pools were determined from water extracts (soil-MilliQ water-suspension = 1:5) that were shaken for 2 h at 150 rpm and filtered (1-2 µm, Ahlström-Munksjö Munktell quantitative filter paper). Soil nitrate and ammonium contents were measured colorimetrically (QuAAtro39 Continuous Segmented Flow Analyzer, SEAL analytical, Norderstedt, Germany). Soil total dissolved N (TDN) concentrations were based on the N concentrations determined by Isotope Ratio Mass Spectrometry (Flash EA 2000 coupled to a DeltaV IRMS, Thermo Fisher Scientific, Bremen, Germany), except for March and August 2020 where all samples were determined with a TOC/TN analyzer (Formacs^HT-I^ TOC/TN Analyzer, Skalar Analytical B.V., Breda, The Netherlands).

### DNA extraction and quantitative PCR

Soil samples were freeze-dried, then the two replicate cores per time point per plot were pooled and milled, resulting in a total of 150 soil samples. A subsample of 50 mg of the soil powder was used for DNA extraction (NucleoSpin Soil kit, Macherey-Nagel, Düren, Germany) with buffer SL1 (without Enhancer SX) according to the manufacturer’s instructions.

Total archaeal, bacterial and fungal abundances were determined via quantitative PCR (qPCR) using the same primer pairs as used for the sequencing (see below; Table S1). In addition, the genetic potential for N_2_ fixation (*nifH*), ammonia oxidation (archaeal and bacterial *amoA*), nitrite reduction in the denitrification pathway (*nirK* and *nirS*), nitrous oxide reduction (*nosZI* and *nosZII*), and nitrite reduction via dissimilatory nitrate reduction to ammonium (*nrfA*) was quantified by qPCR. All qPCR reactions contained IQ 1× SYBR® Green Supermix (BIO-RAD, CA, USA), 0.1% bovine serum albumin (BSA; New England Biolabs, Ipswich, MA, USA; not added to archaeal 16S rRNA gene assays), primers, water and 4-15 ng of template DNA, and were performed on a BIORAD CFX Connect Real-Time system (BIO-RAD). To check for PCR inhibition, all samples and non-template controls were spiked with a known amount of pCR4-TOPO (Thermo Fisher Scientific, Waltham, MA, USA) and amplified using plasmid-specific M13F and M13R primers. No inhibition was detected with the amount of DNA used. Standard curves were obtained by serial dilutions of linearized plasmids of each amplified gene fragment. Amplicons were checked for length and specificity by qPCR melting curve analysis and by agarose gel electrophoresis. Data is expressed as copy numbers per g dry weight (DW) soil. Detailed information on primers, cycling conditions, and efficiencies can be found in Table S1.

### PCR and sequencing

The V4-V5 hypervariable region of the bacterial 16S rRNA gene was amplified with primer pair 515F (Parada et al., 2016) and 926R (Quince et al., 2011). The V3-V4 hypervariable region of the archaeal 16S rRNA gene was amplified with primer pair S-D-Arch-0349-a-S-17 (Takai & Horikoshi, 2000) and S-D-Bact-0785-a-A-21 (Herlemann et al., 2011). Each sample was amplified in duplicate 15 µL reactions containing 1× Phusion High-Fidelity PCR Master Mix (Thermo Fisher Scientific), 0.25 µM of each primer, 0.1% BSA, water, and template DNA. After the first PCR, the duplicates were pooled, checked by gel electrophoresis, and cleaned with Sera-Mag beads (Merck KGaA, Darmstadt, Germany). In a second PCR with similar conditions, the cleaned PCR 1 products were amplified again with primers (0.2 µM) containing the sample-specific sequencing index combination for 8 cycles at an annealing temperature of 55°C. Details on cycling conditions of PCR 1 and 2 can be found in Table S2. The final PCR products were checked via gel electrophoresis, cleaned with Sera-Mag beads, and their concentration was measured with a Qubit fluorometer (Thermo Fisher Scientific). Amplicons from all samples were pooled in equimolar amounts and the two resulting libraries (one bacterial and one archaeal) were cleaned and concentrated. The libraries were quality checked on a 2100 BioAnalyzer (Agilent, Santa Clara, CA, USA) and sequenced on an Illumina MiSeq platform (2×250 bp) by SciLifeLab (Uppsala, Sweden). The bacterial library was sequenced using two flow cells to ensure sufficient sequencing depth.

Fungal communities were amplified with primers gITS7 (Ihrmark et al., 2012) and ITS4 (White et al., 1990)/ITS4A (Sterkenburg et al., 2018), each containing a sample-specific identification tag (Clemmensen et al., 2023). The 50 µL reactions contained 0.025 U Dream Taq polymerase (Thermo Fisher Scientific), 1× buffer, 200 µM dNTPS, 0.75 mM MgCl_2_, tagged primer mix, water, and template DNA. Details on cycling conditions can be found in Table S2. To avoid overamplification and length-based biases, PCRs were run with minimized number of cycles (Castaño et al., 2020). PCR products were purified with the AMPure kit (Beckman Coulter, Brea, CA, USA) and quantified with the Qubit fluorometer. The PCR products were mixed into two pools in equal amounts, and the sequencing pools were further purified with EZNA Cycle Pure kit (Omega Bio-Tek, Norcross, GA, USA) and checked for correct length distribution on a BioAnalyzer 7500 chip (Agilent Technologies, Santa Clara, CA, USA). Each pool was sequenced on a Pacific Biosciences Sequel 1 SMRT cell after addition of sequencing adapters by ligation at Uppsala Genome Center (SciLifeLab, Uppsala, Sweden).

### Sequence analysis

Sequence analysis for bacterial and archaeal amplicons was performed using the R software v. 4.2.2 (R Core Team, 2021). Amplicon sequence variants (ASVs) were inferred using the ‘dada2’ workflow v. 1.26.0 (Callahan et al., 2016). Remaining primer sequences after the filter and trim step were removed with Cutadapt v. 4.2 (Martin, 2011). Chimeras were discarded using the function *removeBimeraDenovo* and the resulting ASVs were classified against the SILVA database v. 138.1 (Quast et al., 2012). ASVs classified as ‘Chloroplast’ or ‘Mitochondria’ and singletons were discarded and the non-bacterial and non-archaeal sequences were excluded for the bacteria and archaea datasets, respectively. The resulting bacterial dataset consisted of 29,466 ASVs and 7,174,779 reads and the archaeal dataset consisted of 1068 ASVs and 2,003,096 reads.

Sequences for fungal community analysis were quality filtered and clustered using the bioinformatics pipeline SCATA (http://scata.mykopat.slu.se/; Ihrmark et al., 2012). High quality sequences (>20 in average score and >3 of individual bases), over 140 bp long and with 100% match with the sample-identification tags and at least 90% match with the primer sequences passed quality control. After removing sequences only found once, the remaining sequences were clustered into species level clusters by single-linkage clustering using a 98.5% sequence similarity criterion for pairwise comparisons (Usearch; Edgar, 2010). All fungal reference sequences in the UNITE database (https://unite.ut.ee/; Abarenkov et al., 2010) were included passively in the clustering to verify species delimitation and provide identification. Clusters thus simulate species hypotheses as defined in UNITE (Koljalg et al., 2020), and we hereafter refer to fungal clusters as “species”. After excluding clusters with less than 4 reads and non-fungal clusters (74; 73996 reads), the resulting dataset consisted of 1161 fungal species and 128,278 reads. Next, the 380 fungal species with at least 40 reads (95% of total reads) were further assessed for taxonomic identity and functional guild, by comparing reference sequences to annotated UNITE and INSD reference databases using the massBLASTer and SHmatching tools in the UNITE PlutoF module. At least 98% similarity was required for species identification, otherwise species were identified to higher taxonomic levels or left unidentified. The following fungal guilds were classified: ectomycorrhizal (ECM) fungi, other root associated fungi (including ericoid mycorrhizal species) and saprotrophic fungi with separate subgroups of molds and yeasts. By combining the relative proportion of fungal guilds from sequencing and total fungal abundances determined by qPCR (see above), we estimated absolute abundances of each fungal guild.

Rarefaction curves were generated with the ‘vegan’ package v. 2.6.4 (Oksanen et al., 2022) to check for sufficient sequencing depth and samples without asymptotic curves were filtered out. Next, the ASV and species tables were rarefied using the function *rrarefy* and split into frequent and rare ASVs based on the species abundance model (Hubbell, 2011) to reduce the influence from rare taxa. The frequency of occurrence (Magurran & Henderson, 2003) was then used to model whether ASVs/species followed a stochastic (Poisson) distribution and those falling below the 2.5 % confidence limit of the χ^2^ distribution were discarded (Krebs, 1999). The final datasets contained 1411 (bacteria), 131 (fungi) and 371 (archaea) frequent ASVs/species.

### Statistical analysis

Temporal effects on microbial communities were assessed by two complementary approaches. To test for temporal variation among all single time points over the year, sampling time points were treated as individual levels. To evaluate seasonal patterns, we grouped time points into seasons as defined in Table 1. To account for potential interannual variability, time points grouped as summer in 2019 and 2020 were treated as separate seasonal categories (summer1 and summer2). This approach avoids assuming recurring seasonal patterns and allows differences between years to be explicitly evaluated. All statistical analyses were conducted in R v. 4.2.2 (R Core Team, 2021). To test for significant differences in microbial abundances (qPCR) between sampling sites and time points or seasons, we used linear mixed-effects models (LMEM) with site and either time point or season as fixed effect, while block nested within site was included as a random effect (‘nlme’ package, v. 3.1-161, (Pinheiro & Bates, 2000). For this, gene abundances were log_10_-transformed, and model assumptions of normality and homoscedasticity were visually assessed using QQ-plots and fitted against residual plots.

**Table 1:**
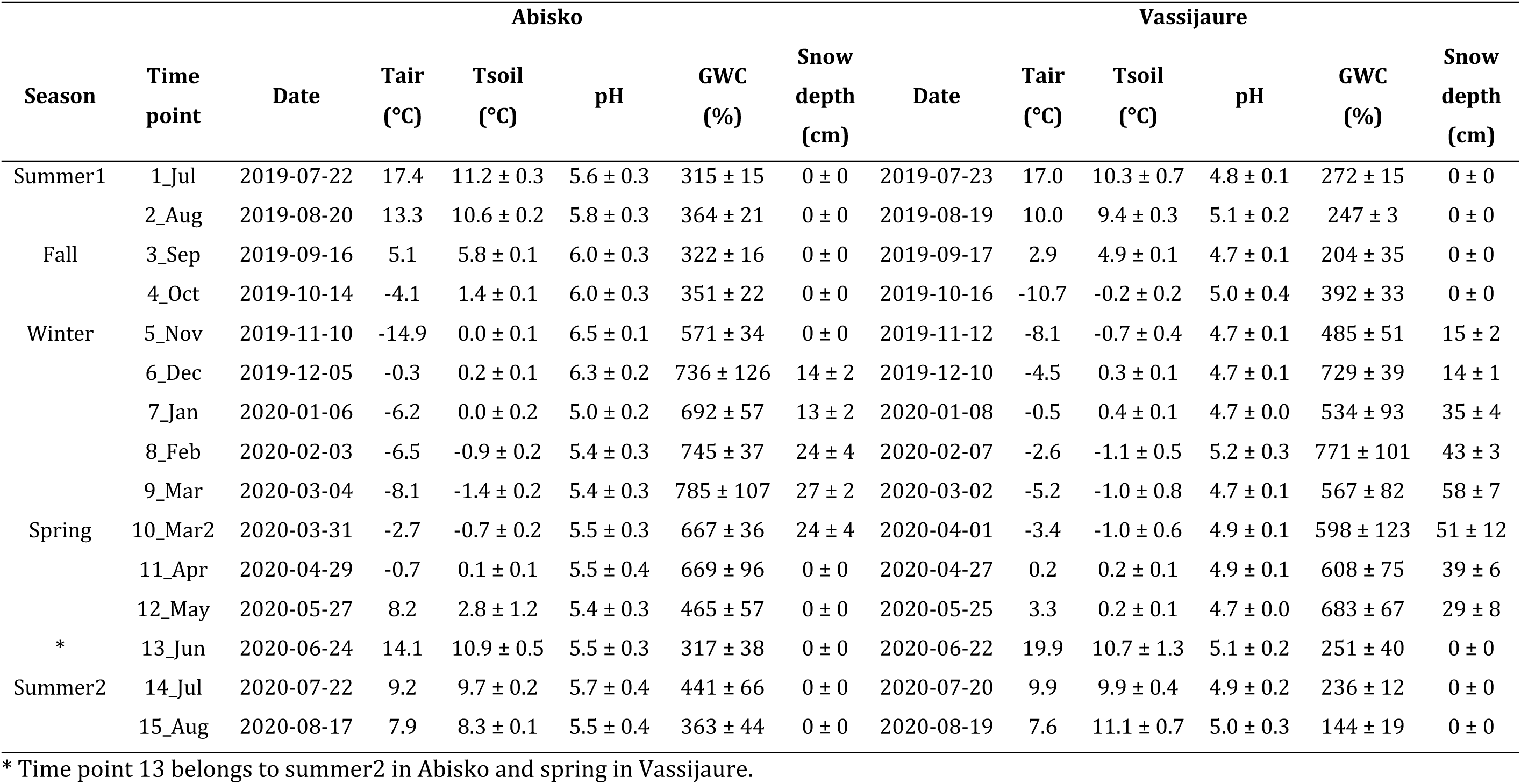
Environmental and soil properties from July 2019 to August 2020 at two sub-Arctic tundra sites in Abisko and Vassijaure as mean values ± standard error (Tair - Air temperature, Tsoil - Soil temperature, GWC - Gravimetric water content).

Correlations between gene abundances and environmental variables, as well as among different microbial groups, were assessed using generalized least squares models (‘nlme’ package). Separate models were fitted for each response–explanatory variable combination, accounting for repeated measurements within blocks using a compound symmetry correlation structure. Model significance was evaluated using t-tests. QQ-plots and fitted vs. residual plots were inspected visually to confirm model assumptions.

Patterns in microbial community composition were visualized using a partial principal component analysis (p-PCA) conditioned to block to account for the spatial structure, and were based on centered log-ratio (clr) transformed relative abundances of ASVs (bacteria, archaea) and species (fungi). Variation in microbial community composition among the two sampling sites was tested using a permutational analysis of variance (PERMANOVA) as implemented in the *adonis* function of the ‘vegan’ package. Site effects were tested using 999 permutations constrained within time points. Next, for each site, the effect of sampling time point and season was tested in separate PERMANOVAs with 999 permutations constrained within blocks. For all PERMANOVAs, the community data were clr-transformed and the PERMANOVA was based on Euclidean distances. Homogeneity of multivariate dispersions was evaluated prior to PERMANOVA. We analyzed the effect of time point and season in two different models.

Time–decay relationships in community composition were examined by relating pairwise community dissimilarity to temporal separation between samples. Community dissimilarity was quantified as Euclidean distance obtained from a p-PCA that accounted for block effects (see above). Temporal distance was calculated as the number of days between sampling dates for each pair of samples. All non-zero pairwise comparisons were retained and the relationship between temporal distance and community dissimilarity was visualized. Further, temporal community turnover was calculated based on the ‘codyn’ package v. 2.0.5 (Hallett et al., 2016). Due to the high spatial variability within each block, the community data was summed across blocks to obtain a site-level community for each time point prior to analysis. Total turnover as well as the proportion of appearing and disappearing taxa were then calculated between pairs of consecutive time points.

To identify the main environmental drivers of microbial community variation while accounting for spatial structure, partial redundancy analysis (p-RDA) was performed using the ‘vegan’ package. First, to evaluate temporal scales, we generated distance-based Moran’s eigenvector maps (MEM) from sampling time points (days) using the ‘adespatial’ package v. 0.3-23 (Dray et al., 2022) to model temporal autocorrelation. Significant MEM variables were identified using forward selection. Second, community composition data were then constrained by selected environmental variables and the significant MEMs, while block was included as a conditioning variable to account for the spatial structure. Model selection was performed using stepwise forward selection to identify the most parsimonious set of explanatory variables.

For the identification of seasonal abundance profiles (bacterial or archaeal ASV and fungal species), we used the ‘timeOmics’ package v. 1.6.0 (Bodein et al., 2021) in R v. 4.1.2 (R Core Team, 2021). To account for block effect, the relative abundances of taxa were clr-transformed per sampling site, and a LMEM was fitted with block as random effect. Residuals were then used as input data for the seasonal cluster analysis. If the model failed for a taxon, a simple linear model was used instead. Next, we fitted a penalized-spline mixed-effects model with function *lmmspline* and basis = “p-spline” from the ‘lmms’ package v. 1.3.3 (Straube et al., 2016) to the 15 time points and retained only well-modelled curves with *wrapper.filter.splines*. The spline-fitted values were then subjected to dimension reduction and clustering by principal component analysis using function *pca* from the ‘mixOmics’ package v. 6.18.1 (Rohart et al., 2017); the optimal number of clusters/components was chosen by inspecting plotIndiv, plotVar and plotLoadings. Finally, seasonal clusters and their member taxa were visualized and exported with plot.Long, providing clusters of ASVs/species that share coherent seasonal trajectories. An approximate maximum-likelihood phylogenetic tree of seasonal bacterial and archaeal ASVs was inferred using FastTree under a GTR model based on aligned sequences (SINA 1.7.2.; Pruesse et al., 2012)). The resulting phylogenetic tree was visualized with iTOL v. 7.5 (Letunic & Bork, 2007).

## Results

### Environmental conditions

The daily mean soil temperature ranged from −1.4 to 11.2°C in Abisko and from −1.1 to 11.1°C in Vassijaure (Table 1). Most freeze-thaw cycles occurred in October till December at both sites, while additional, less frequent zero-degree transitions occurred in January and February in Vassijaure in the blocks with the highest snow-cover, allowing relatively stable, high soil temperatures (Fig. S2). The onset of the snow-covered period was nearly at the same time at both sites, in November (Vassijaure) and December (Abisko), whereas the snow melt occurred two months earlier in Abisko (April) than in Vassijaure (June) (Table 1). The maximum snow cover at Vassijaure (58 ± 7 cm, mean ± standard error) was approximately twice as deep as in Abisko (27 ± 2 cm). The soil moisture (based on thawed soil samples) was highest from November till May at both sites and may not reflect water availability in the field when soils were frozen. Soil pH was on average higher in Abisko (5.7 ± 0.1) than in Vassijaure (4.9 ± 0.0). While the soil pH remained relatively stable over the one-year time-period at Vassijaure, it dropped from December to January in Abisko.

### Soil N pools

Soil TDN content was similar at both sites (Fig. 1) and peaked in early and late winter, except for lower levels in January and February. Soil ammonium and nitrate contents were generally low or below detection limit with some higher values from October to February and in July 2020 (ammonium) and in July 2019 and from October to December (nitrate).

**Figure 1:**
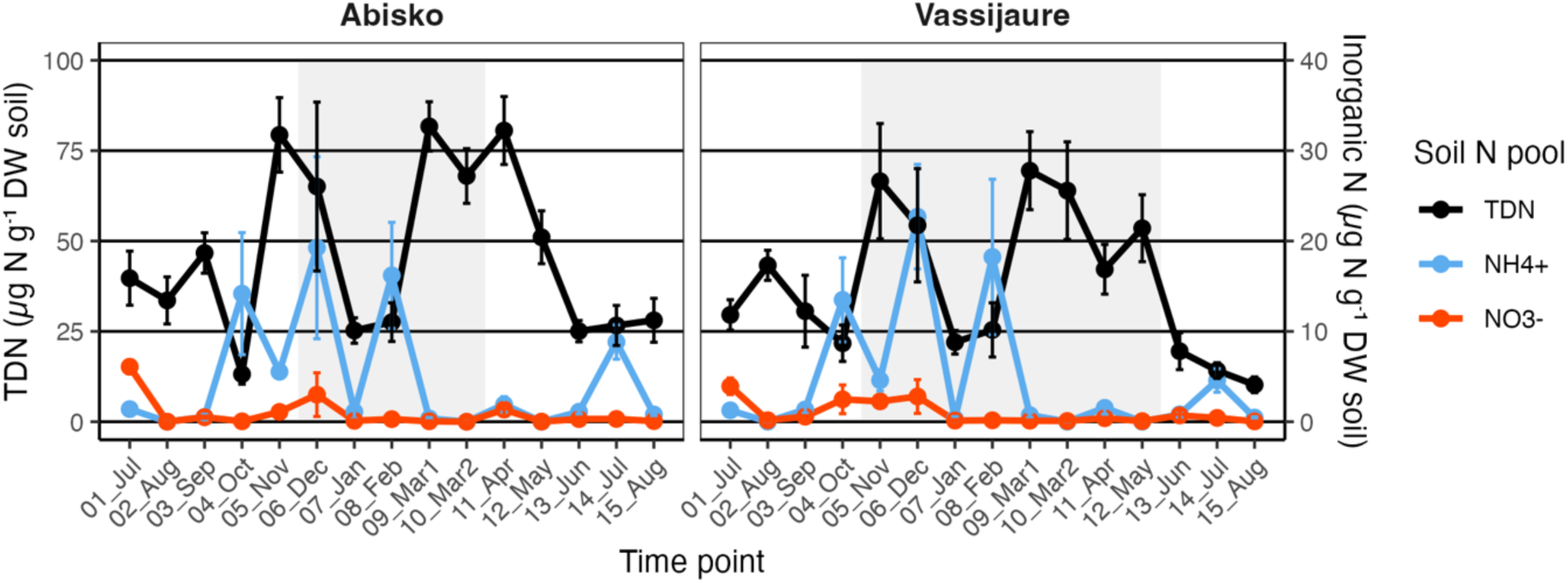
Temporal variation in soil nitrogen pools. (TDN, ammonium, and nitrate concentrations) during one year from July 2019 to August 2020 at two tundra sites in Abisko and Vassijaure. Concentrations are plotted as means and error bars denote standard errors (n = 5). Values below detection limit (DL) were converted to DL:2 before averaging and accounted for 15.3% and 25.3% of all samples (n = 150) for ammonium and nitrate, respectively. The grey-shaded background corresponds to the duration of the snow-covered period.

### Abundances of taxonomical groups

In both tundra sites, abundances of bacteria, fungi, and archaea fluctuated significantly over the year and overall trends for each group were similar between the sites (Fig. 2, Table 2). Both bacteria and fungi increased towards winter, with increasing relative dominance of fungi in fall and early winter and bacteria in late winter (as reflected by the fungi:bacteria ratio). Archaea showed the opposite pattern with lowest abundances in early winter. However, in Abisko, fungal abundances dropped by almost one order of magnitude in December co-occurring with a decline in archaeal abundances, while bacterial abundances only decreased slightly. Bacterial abundances did not differ significantly between the sites, whereas fungal and archaeal abundances were higher in Vassijaure (Fig. 2, Table 2). Bacterial and fungal abundances were correlated at both sites (Fig. 3) with a stronger synchrony at Vassijaure as compared to Abisko (Fig. 2). Bacterial abundances were also correlated to archaeal abundances in Vassijaure.

**Figure 2:**
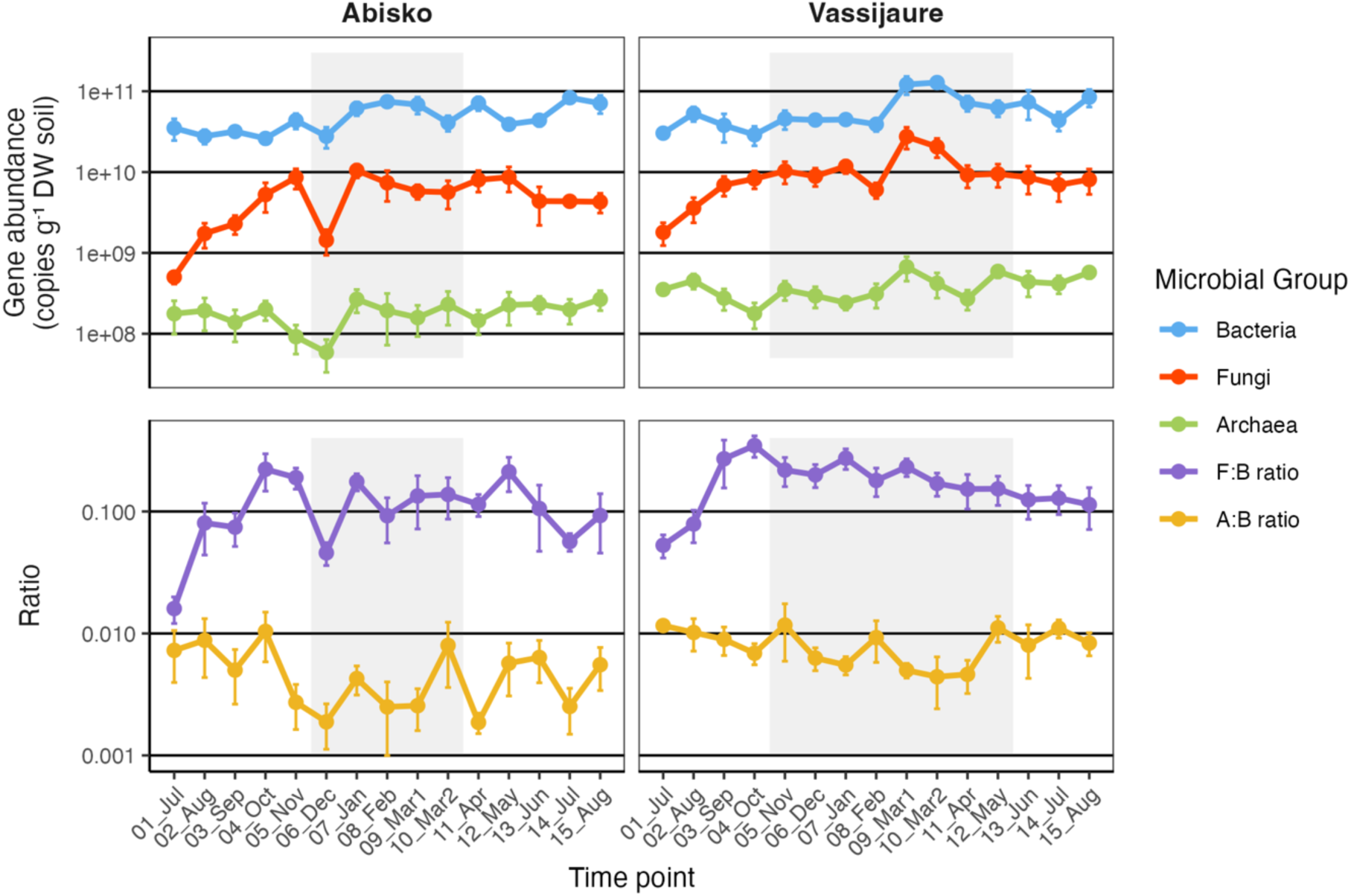
Temporal variation in gene copy numbers for microbial taxonomic groups (Bacteria, Fungi, and Archaea) and their ratios during one year from July 2019 to August 2020 at two tundra sites in Abisko and Vassijaure. Gene abundances and ratios are plotted on a log_10_ scale and show means with error bars denoting standard errors (n = 5). The grey-shaded background corresponds to the duration of the snow-covered period. F:B ratio – fungi:bacteria ratio, A:B ratio – archaea:bacteria ratio.

**Figure 3:**
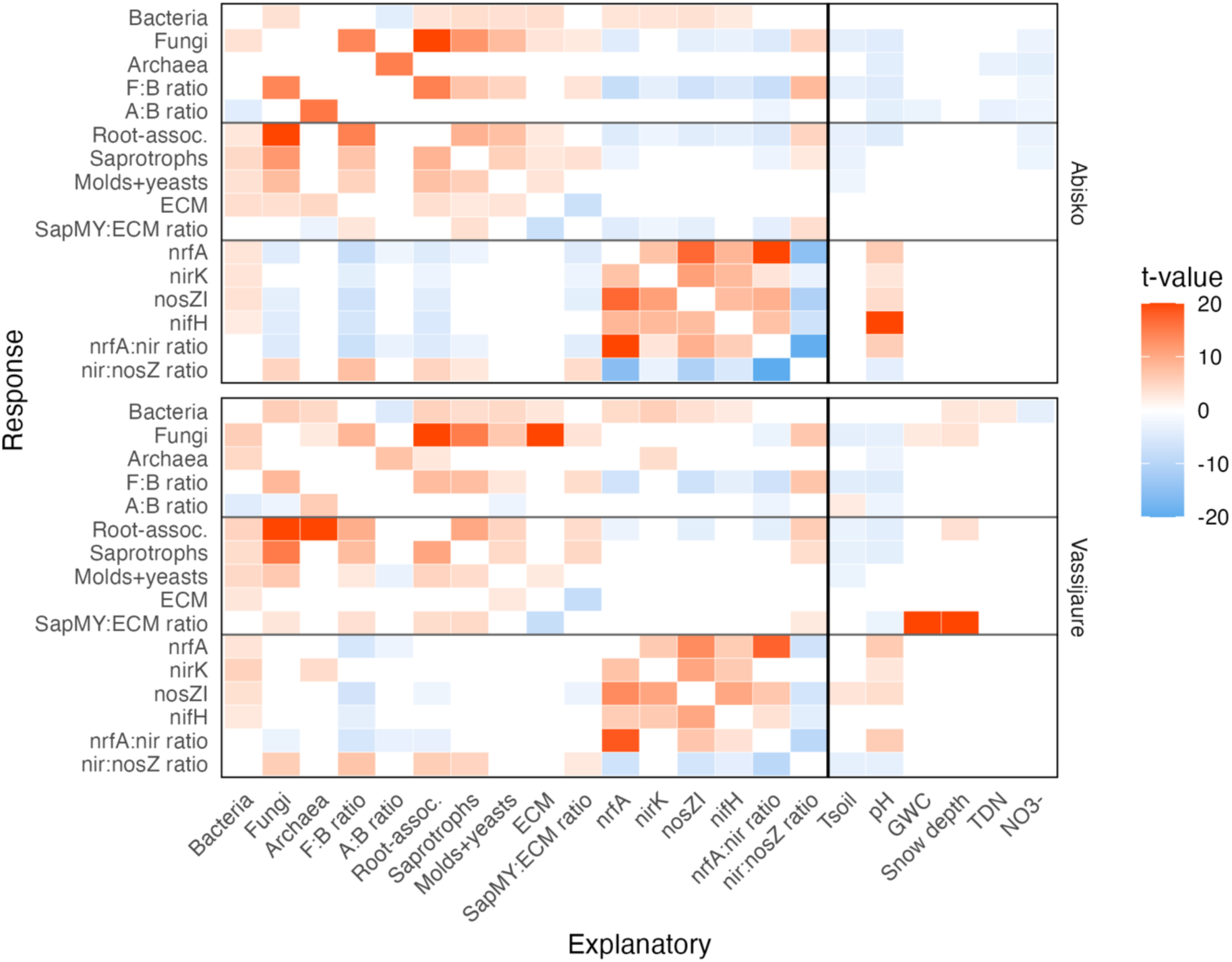
Heatmap of associations among microbial groups and between microbial groups and abiotic variables for the two tundra sites. Tiles show t-values from generalized least squares (GLS) models accounting for repeated measurements within blocks over time. Positive associations are shown in red and negative associations in blue; non-significant associations after false discovery rate (FDR) correction (p > 0.05) are shown in white. The color scale for t-values was limited to the range −20 to 20 to improve visual contrast; values outside this range are displayed at the scale limits. Vertical lines separate biotic predictors from abiotic predictors, and horizontal lines indicate the main microbial groups (taxonomic groups, fungal guilds, and functional genes). Panels show results for each site. F:B ratio – fungi:bacteria ratio, A:B ratio – archaea:bacteria ratio, SapMY:ECM ratio – (saprotrophs+molds+yeast):ECM fungi ratio.

**Table 2:**
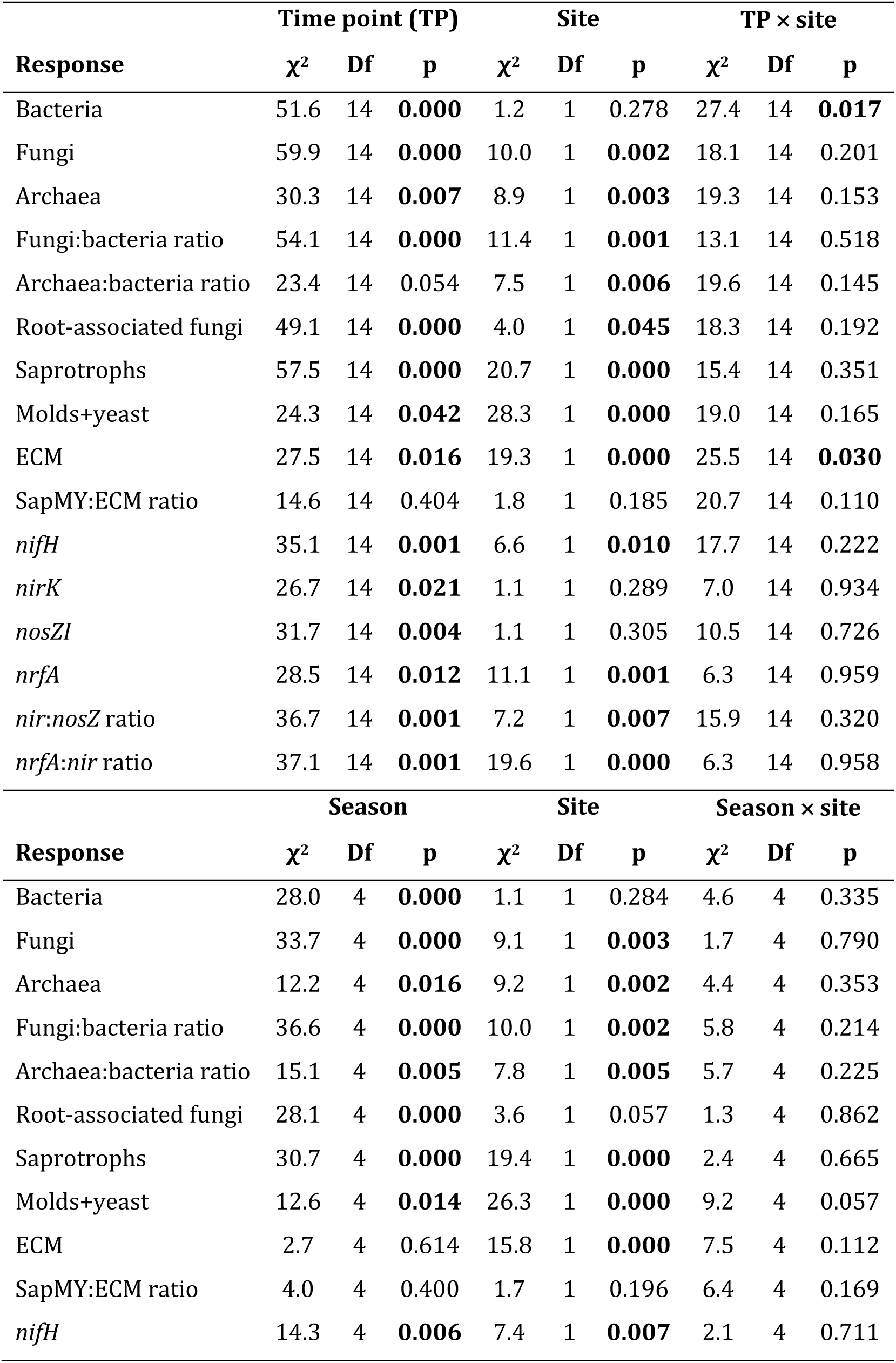
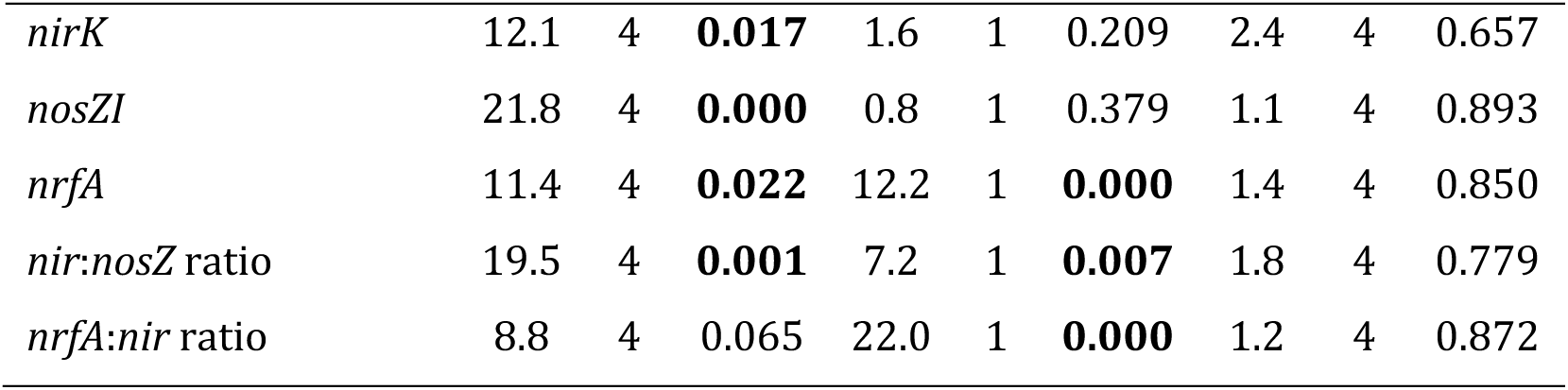
Effect of time point, sampling site and their interaction (two-way ANOVA), and season, sampling site and their interaction (two-way ANOVA) on microbial abundances and ratios. All response variables were log_10_-transformed. Effects with p < 0.05 are shown in bold. SapMY:ECM ratio – (saprotrophs+molds+yeast):ECM fungi ratio.

Bacterial abundances were positively correlated with soil TDN and negatively with nitrate contents in Vassijaure, whereas fungal abundances were negatively related to soil pH and soil temperature at both sites (Fig. 3). In addition, bacterial and fungal abundances were positively linked to snow depth only at the Vassijaure site. Archaeal abundances were negatively related to soil pH at both sites and to soil TDN and nitrate contents only in Abisko.

### Community composition and turnover

The analysis of bacterial and archaeal community composition revealed significant temporal differences among the sampling time points at both sites despite a high within-site spatial variability (Fig. 4, Table S3). In comparison, fungal composition seemed to be more stable across the time points. However, the community composition of all three groups varied also significantly with season and between the Abisko and Vassijaure sites (Fig. 4, Fig. S3, Table S3). The temporal changes in community composition were reflected by increasing Euclidean distances with increasing time spans between sampling time points, although this was less pronounced for fungal communities as compared to bacterial and archaeal communities (Fig. S4). However, both bacterial and fungal community dissimilarity were lower for the longest time span, especially at Vassijaure. This suggests that the summer communities in 2019 and 2020 were more similar to each other than summers were to winters, indicating temporal dynamics were cyclic, while the opposite was true for Archaea.

**Figure 4:**
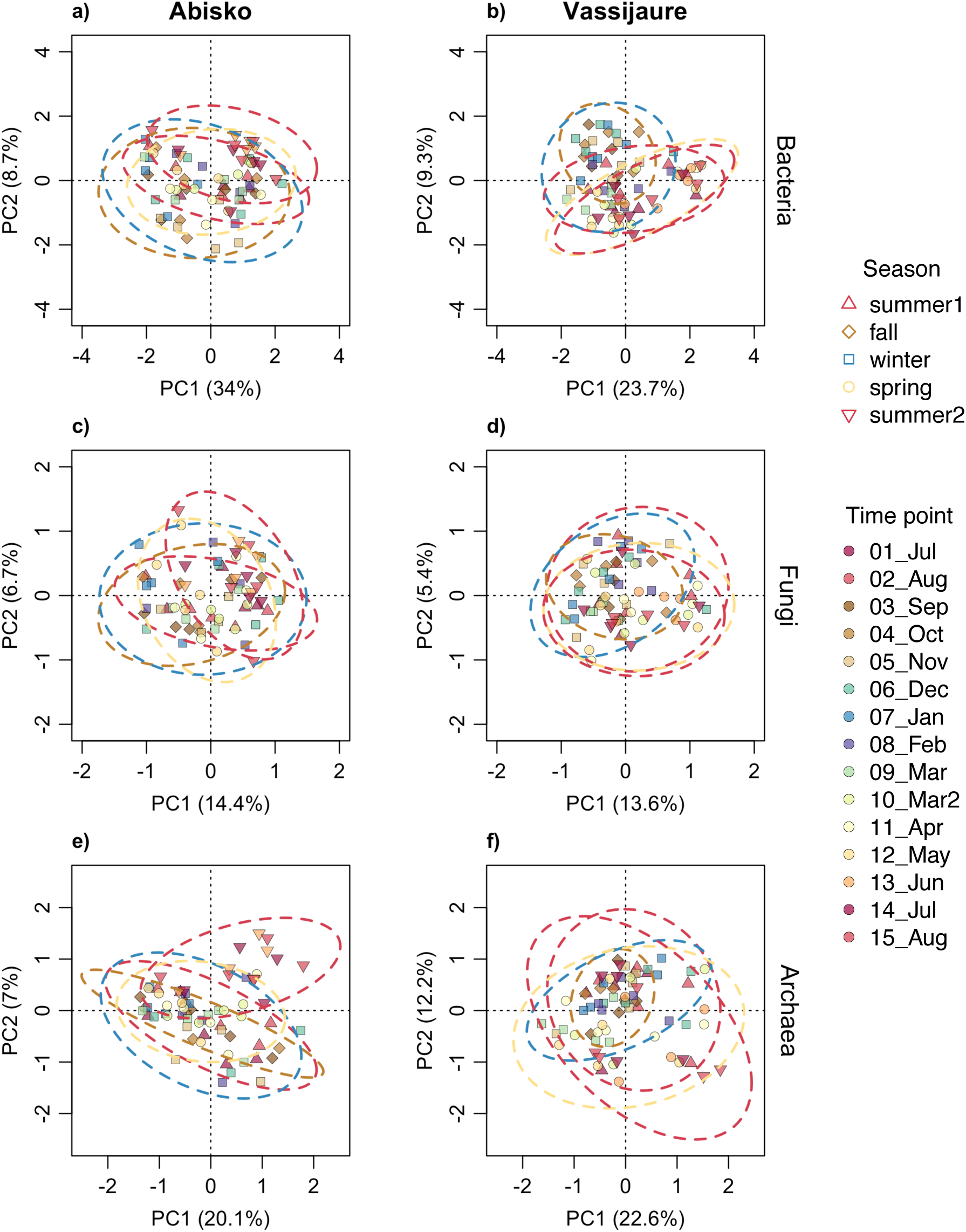
Partial principal component analysis (p-PCA) conditioned for block for bacterial (a, b), fungal (c, d), and archaeal (e, f) communities based on clr-transformed relative ASV/species abundances during one year from July 2019 to August 2020 at two tundra sites in Abisko and Vassijaure. Symbol colors denote time point and symbol shapes denote season. Ellipses indicate within-season dispersion in PCA space, scaled to encompass approximately 95% of the variability.

Temporal variation in community structure of all microbial groups was correlated with the broadest temporal scale (MEM1) in most cases, shifting during the winter months (Table S4, Fig. S5). Only archaeal community structure at Abisko was correlated to finer temporal scales (MEM2, MEM4). When these significant temporal vectors (MEMs) together with the other environmental variables were included in a partial RDA, forward selection revealed that the variation in all three microbial communities was strongly linked to firstly soil pH and secondly MEM1 (except for archaea at Abisko: MEM2 + MEM4) at both sites (Table S5). Bacterial community composition was also associated with soil temperature at Vassijaure. Further, in Abisko, snow depth explained a significant proportion of the variation of archaeal communities.

The temporal patterns of total community turnover calculated as the summed proportion of appearing and disappearing ASVs or fungal species relative to all present for two consecutive time points varied between the two sampling sites (Fig. 5). Surprisingly, community turnover of all three taxonomical groups peaked in mid-winter. In Abisko, this mid-winter peak occurred in January or February and there was a second smaller peak in fall (fungi) or spring (bacteria and archaea). In Vassijaure, the mid-winter peak was delayed by one month (February or March) and was less pronounced for bacteria and archaea, while fungi had highest total turnover in summer (August 2019). The mid-winter peak was mainly driven by appearing ASVs for bacteria, while both appearing and disappearing ASVs caused the mid-winter peak for archaea.

**Figure 5:**
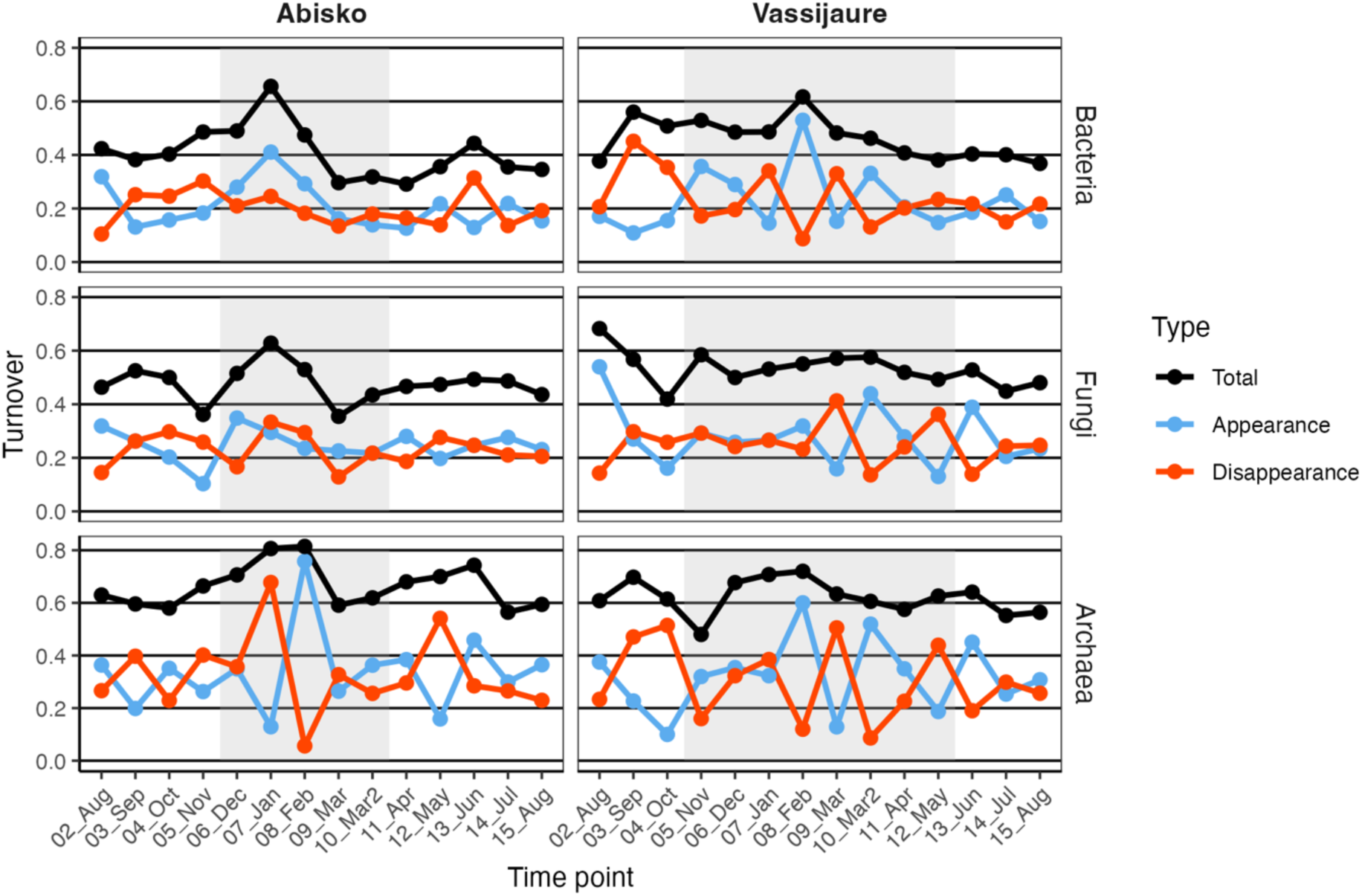
Variation in microbial community turnover (total, appearance, disappearance) for bacteria, fungi, and archaea during one year from July 2019 to August 2020 at two tundra sites in Abisko and Vassijaure. Turnover represents the proportion of appearing, disappearing or both (total) taxa in relation to the sum of all taxa between two time points. Turnover was calculated based on sums of ASVs/species across all five blocks. The grey-shaded background corresponds to the duration of the snow-covered period.

### Seasonal synchronous taxonomic clusters

All three microbial groups had subsets of seasonal taxa that were either more abundant in winter or in summer (Fig. 6). The bacterial clusters included 23 to 69 ASVs, the fungal clusters consisted of 1 or 2 species, and the archaeal clusters consisted of 4 or 5 ASVs (Table S6). Bacterial seasonal ASVs were diverse and were affiliated with 14 phyla and 42 different orders (Fig. S6, Table S6). Some phyla contained predominantly either summer or winter clusters, for example most seasonal Actinobacteriota ASVs were more abundant in summer, while most seasonal Planctomycetota and Bacteroidota ASVs dominated in winter. For other phyla, there was a seasonal signal on a lower taxonomic level such as order, e.g. the Proteobacteria order Acetobacterales was more dominant in winter, whereas most ASVs affiliated with the order Rhizobiales were dominant in summer. Bacterial winter ASVs, that were detected in both sites, belonged to the orders Gammaproteobacteria Incertae Sedis, Acetobacterales, Acidobacteriales, and Chthonomonadales. Bacterial summer ASVs from both sites belonged to the orders Myxococcales (n=2), Acidobacteriota Subgroup 17, IMCC26256 (Actinobacteriota), Solirubrobacterales, and Frankiales.

**Figure 6:**
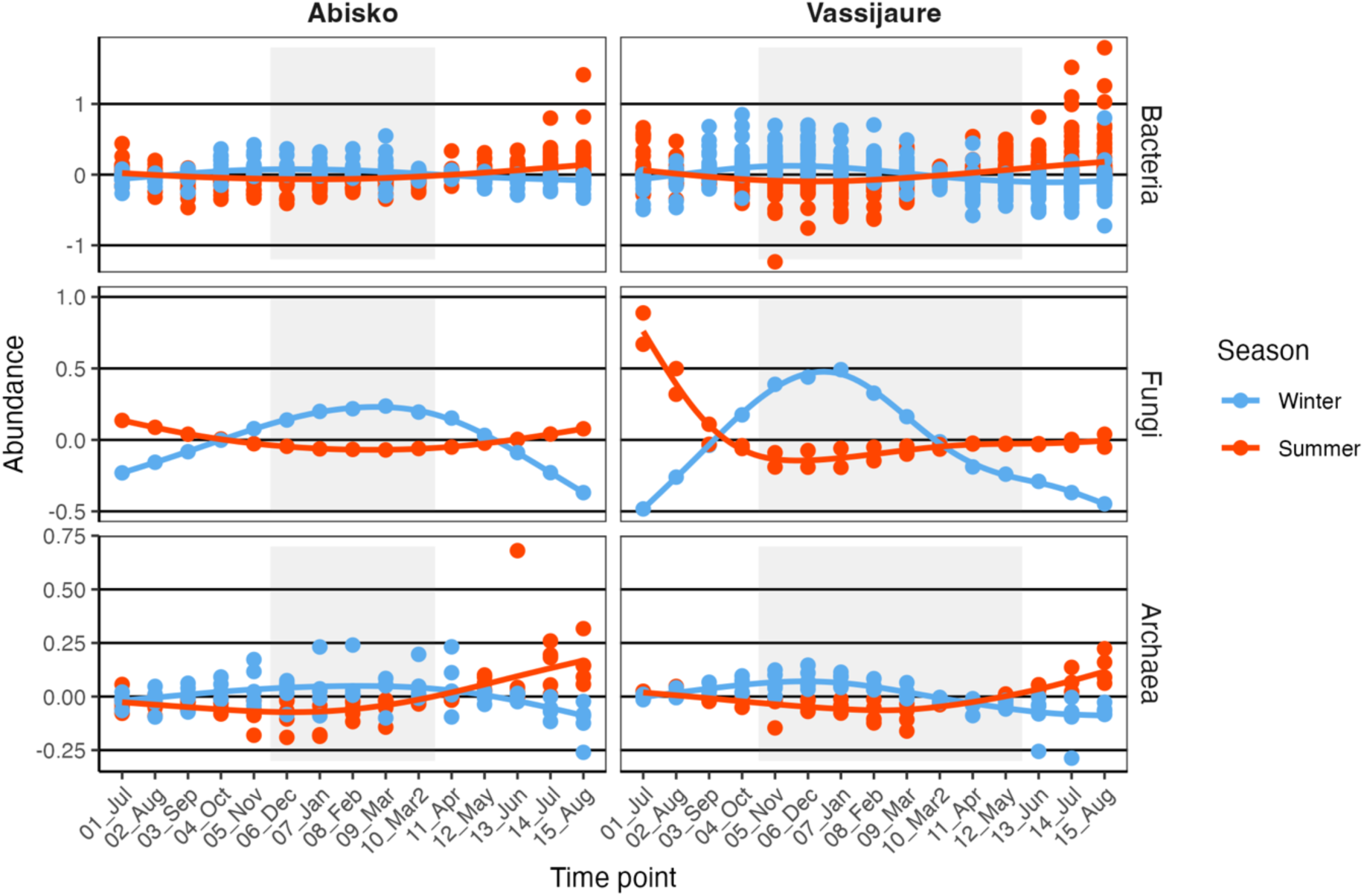
Variation in abundances of seasonal taxa for bacteria, fungi, and archaea during one year from July 2019 to August 2020 at the two sampling sites Abisko and Vassijaure. Note that one outlier (abundance = 1.1) is not shown for the Archaea winter cluster at Vassijaure in September. The grey-shaded background corresponds to the duration of the snow-covered period.

The fungal winter species were affiliated with Herpotrichiellaceae and *Hyaloscypha sp*. in Abisko and Vassijaure (Table S6), respectively, both of which belonged to the putative group of root-associated ascomycetes. Fungal summer species both belonged to the ectomycorrhizal *Russula* genus (*R. purpureofusca* in Abisko and *R. renidens* in Vassijaure). An additional fungal summer species belonged to the class Sordariomycetes (mainly litter saprotrophs) and was only detected in Vassijaure.

The archaeal winter ASVs belonged to the orders Group 1.1c, Woesearchaeales, or Micrarchaeales (only in Abisko) and included one Group 1.1c ASV, which was present at both sites (Fig. S6, Table S6). In contrast, the summer clusters were site-specific and belonged either to Nitrososphaerales and Woesearchaeales (Abisko) or Group 1.1c and Marine Group II (Vassijaure).

### Abundances of fungal guilds

The abundances of root-associated fungi and saprotrophs increased over fall and winter (Fig. 7) following the trend of total fungal abundances (Fig. 2). In contrast, ECM fungi and molds+yeasts appeared to vary less during the one-year period due to a larger variation among replicates (Table 2). Noticeably, all fungal guilds showed a steep decline in December in Abisko with ECM fungi decreasing the most by a factor of approximately 60 while root-associated fungi, molds+yeasts, and other saprotrophs decreased by a factor of approximately 7, 5.5, and 3.5, respectively. Overall, all fungal guilds showed on average higher abundances in Vassijaure than in Abisko. Although the SapMY:ECM ratio (saprotrophs+molds+yeast:ECM fungi), a proxy for the relative importance of N mineralization vs. N uptake potential, did not vary significantly between time points or seasons, it was lower in October, January, and April at both sites, indicating reduced potential for inorganic N release by fungi during these months.

**Figure 7:**
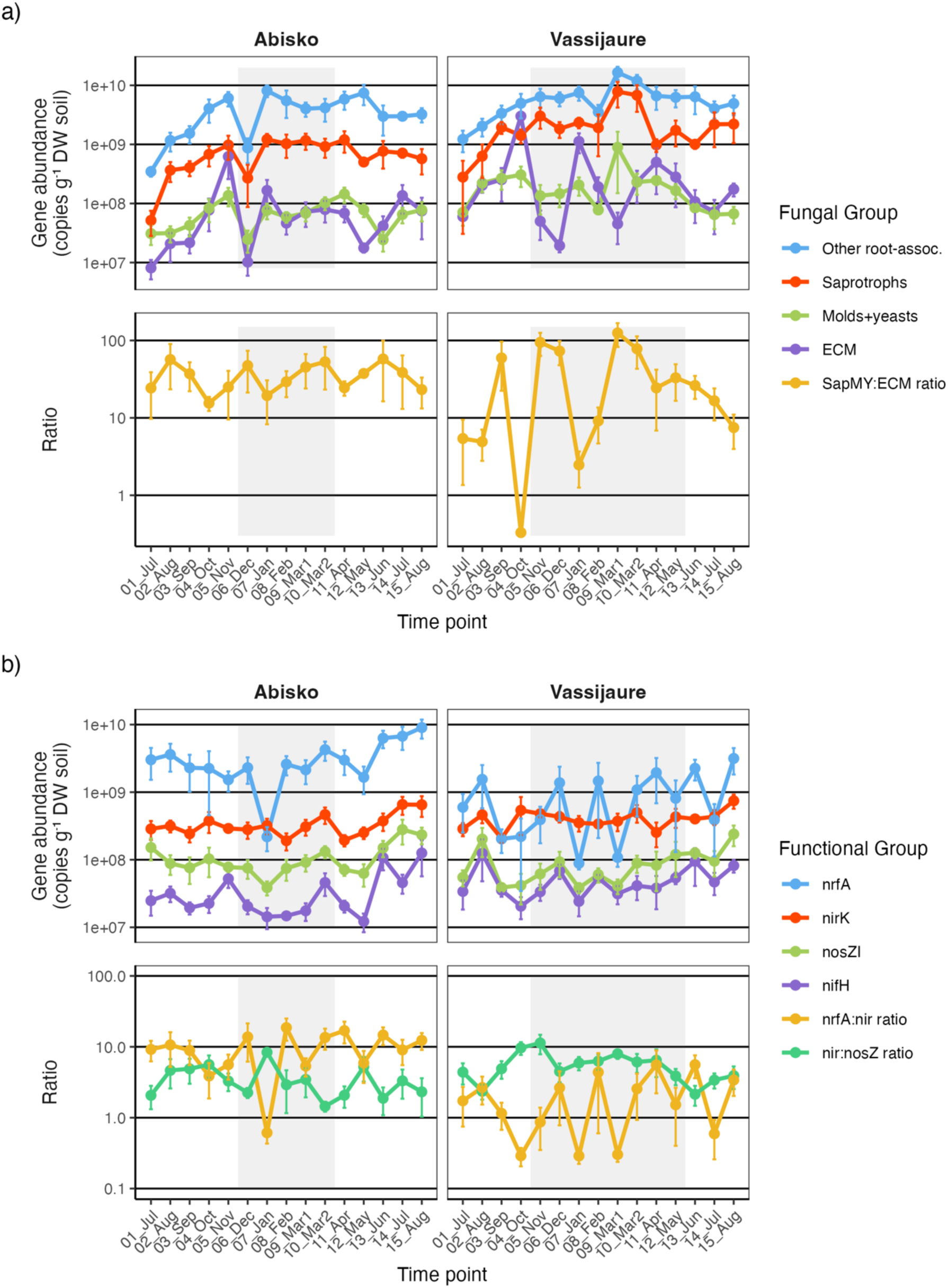
Variation in gene copy numbers and ratios for a) fungal guilds and b) functional N genes during one year from July 2019 to August 2020 at the two sampling sites Abisko and Vassijaure. Gene abundances and ratios are plotted on a log_10_ scale and show means with error bars denoting standard errors (n = 5). SapMY:ECM ratio – (saprotrophs+molds+yeast):ECM fungi ratio.

At both sites, abundances of root-associated fungi, saprotrophs and molds+yeasts were negatively correlated with soil temperature (Fig. 3). Root-associated fungi were negatively correlated with soil pH at both sites, while saprotrophs showed that link only at Vassijaure. Further, in Vassijaure, root-associated fungal abundances and the SapMY:ECM ratio were positively linked to snow depth and the SapMY:ECM ratio was additionally positively linked to soil water content and negatively to soil pH. In contrast, root-associated and saprotrophs fungal abundances were negatively correlated with soil nitrate content in Abisko.

### Abundances of functional N genes

The abundances of all functional N genes varied significantly throughout the year (Fig. 7, Table 2), however the overall trend differed from bacterial abundances. The genetic potentials for nitrite reduction (dominated by *nirK*), nitrous oxide reduction (dominated by *nosZI*), and nitrate ammonification (*nrfA*) were lower in winter and increased towards the second summer. The genetic potential for N_2_ fixation (*nifH*) showed a similar trend with highest abundances in summer (except for the first summer in Abisko). The *nir*:*nosZ* ratio, a proxy for gaseous N loss in the form of the greenhouse gas N_2_O vs. N_2_ during denitrification, peaked in early winter, while the *nrfA*:*nir* ratio, a proxy for the genetic potential for N retention vs. loss was higher in late winter. Ammonia oxidation genes (*amoA*) were dominated by archaea, with a tendency to be present mainly during summer and spring, whereas they were absent or below detection limit in most samples from the snow-covered period (Fig. S7). Abundances of the nitrous oxide reducer from clade II (*nosZII*) had a similar trend as for clade I (*nosZI*) but were only detected in 53% of the samples. Similarly, abundances for *nirS*-type nitrite reducers were only present in 19% of the samples, predominantly in Abisko. Other site-specific differences were found for the abundance of *nifH* and the *nir*:*nosZ* ratio, with higher values at Vassijaure, whereas *nrfA* abundances and the *nrfA*:*nir* ratio were higher at Abisko (Fig. 7). The abundances for *nirK* and *nosZI* did not differ significantly between the two sites (Table 2).

Most functional N gene abundances (*nirK*, *nosZI*, *nrfA*) were positively correlated with soil pH at both sampling sites, while *nifH* was correlated with soil pH only in Abisko (Fig. 3). Abundances of *nosZI* were positively, and the *nir*:*nosZ* ratio was negatively, linked to soil temperature in Vassijaure. Ratios of *nir*:*nosZ* and *nrfA*:*nir* were also negatively and positively linked to soil pH, respectively, at both sites.

### Synchrony of func+onal groups

Abundances of most fungal guilds were positively correlated with each other, with less significant links in Vassijaure, in particular for ECM fungi (Fig. 3). All functional N gene abundances were positively correlated with each other at both sites. The strongest links were found between *nosZI* and *nrfA* (both sites), *nosZI* and *nirK* (both sites), and *nosZI* and *nifH* (Vassijaure). In contrast, some fungal guild abundances were negatively correlated with functional N gene abundances. These correlations included root-associated fungi, saprotrophs, and all functional N genes (predominantly *nrfA* and *nosZI*) with fewer links in Vassijaure.

## Discussion

### Winter microbial communi+es are highly dynamic

Abundances of bacteria and fungi increased towards and during winter (Fig. 2) and community turnover of all three taxonomic groups (bacteria, fungi, and archaea) peaked in mid-winter (except for fungi at Vassijaure) (Fig. 5). These trends indicate that microbial communities in tundra soils are not just surviving but are also growing and highly dynamic during the long Arctic winter. Consequently, our first hypothesis, that microbial abundances decline or remain stable during winter, can be at least partly rejected. Our second hypothesis, that microbial turnover would be highest in spring and fall, was also partly contradicted. However, in line with this hypothesis, bacterial and archaeal communities showed a secondary smaller turnover peak in spring (Abisko) or in early fall (Vassijaure).

Although previous studies reported inconsistent winter trends in microbial abundances (Buckeridge et al., 2013; Schostag et al., 2015), it may seem counterintuitive that abundances peak in winter during sub-zero conditions. Yet several processes may enable microbial growth in winter: relatively stable soil temperatures under snow cover as compared to air temperature fluctuations (Table 1), availability of liquid water (Tilston et al., 2010), continuous plant litter decomposition (Bokhorst et al., 2010; Schmidt & Lipson, 2004), freeze-thaw cycles that release labile carbon and nutrients from lysed cells (Schimel & Clein, 1996), and reduced competitive pressure from plants (Larsen et al., 2012). These factors likely create a temporal niche for enhanced growth in fall and winter, which in our sites appears to be first relatively dominated by fungi and later by bacteria as indicated by the fungi:bacteria ratio. As snow accumulates during the winter and insulates the soil, even frost-sensitive microbial groups may thrive. The positive influence of the snow insulation was reflected by the correlation between snow depth and bacterial and fungal abundances at Vassijaure (Fig. 3).

Surprisingly, winter was not only characterized by the highest microbial abundances, but also by the highest community turnover, indicating substantial shifts in community composition particularly between December and January (Abisko) or January and February (Vassijaure) (Fig. 5). This pattern was in line with the significant temporal scales determined via distance-based Moran’s eigenvector maps, which revealed that the largest temporal scale (MEM1), which indicated a compositional shift during the winter months (Fig. S5), significantly shaped microbial community composition. In addition, the shifting point between winter and summer seasonal taxa also occurred in late winter for most microbial groups (Fig. 6). The timing of the highest community turnover coincided with a drop in soil pH (Table 1), changes in soil TDN, nitrate, and ammonium contents (Fig. 1), and most freeze-thaw cycles occurring before the turnover peak (Fig. S2).

In Abisko, the drop in soil pH (January) was preceded by a drastic decline in fungal, but also bacterial and archaeal, abundances (Fig. 2). This decline could have been caused by the early winter temperature decrease below 0°C and associated freeze-thaw cycles (Feng et al., 2007). Soils in Abisko had a lower proportion of moss-covered ground compared to Vassijaure (Fig. S1), making the soil more vulnerable to freezing (Park et al., 2018; Szymański et al., 2022). Further, snow accumulation was slower, reducing insulation and exposing soil microorganisms to subzero temperatures, potentially causing freeze damage. Freeze-induced microbial die-off may have suppressed OM decomposition and ammonification. Ongoing ammonification releases ammonia, which can consume protons and increases soil pH, thus, reduced ammonification likely contributed to the observed decline in soil pH. The following release of labile compounds from lysed cells potentially stimulated new growth of all three microbial groups after the abundance drop (Fig. 2). In Vassijaure, however, soils were insulated earlier by rapidly accumulating snow and by a generally more prominent moss cover, protecting microbial communities from early winter freeze events. Only when soil temperatures reached a secondary minimum later in winter, declined fungal abundances (February), followed by a smaller decrease in soil pH (March).

At both sites, soil pH was the strongest driver of community compositions for all three microbial groups according to partial-RDA (Table S5). Soil pH is commonly found to be one of the main parameters explaining spatial variation of microbial community composition at various scales (Fierer & Jackson, 2006; Větrovský et al., 2019), but for temporal variation including the cold season in Arctic tundra soils earlier results have been inconclusive (Buckeridge et al., 2013; Männistö et al., 2024; Mundra et al., 2015; Schostag et al., 2015). Interestingly, our study indicates the potential role of soil pH in structuring temporal community trends and highlights the bidirectional interaction between soil pH and microbial communities (Philippot et al., 2024). That is, the decline in soil pH was likely caused by a combination of winter conditions, microbial die-off and altered biotic processes, and, in turn, the changing pH might have contributed to the restructuring of winter communities, which was also reflected by the second most influential variable, the largest temporal scale (MEM1). Overall, soil temperature dynamics, freeze exposure, and site-specific soil conditions appear to have driven microbial declines, soil pH drop, and winter community restructuring.

### Seasonal restructuring of nitrogen cycling poten+al

Most fungal guilds were more abundant in winter or during the shoulder seasons than in the summer periods, however this was particularly clear for root-associated fungi and saprotrophs, while ECM fungi and yeasts and molds were more variable within seasons as well (Fig. 7). In support of this pattern, the fungal species significantly affiliated with winter belonged to Herpotrichiellaceae (Eurotiomycetes; Abisko) and *Hyaloscypha* (Leotiomycetes; Vassijaure), and both species were assigned to the root-associated guild based on matching sequences previously isolated from roots (Abarenkov et al., 2010). The fungal species affiliated with summer both belonged to the ectomycorrhizal *Russula* genus (*R. purpureofusca* in Abisko and *R. renidens* in Vassijaure), suggesting that ECM fungi were more dominant during the warm season, although the large fluctuations within this guild could be linked to functional differences and temporal niche partitioning among ECM fungi. These patterns suggest an annually cyclic pattern with somewhat more dominant saprotrophic communities, and potentially continued OM degradation and N mineralization outside the summer months, and relatively more dominant ECM communities during the warm season, as also found in a temperate forest (Voříšková et al., 2014). Seasonal bacterial ASVs that predominantly occurred in fall and early winter included taxa potentially involved in degradation of (complex) OM (e.g. Solibacterales, Tepidisphaerales, Cytophagales, Chitinophagales, Planctomycetota orders) (Fig. S6, Table S6). These findings are consistent with elevated winter exoenzyme activities in an alpine tundra soil (Broadbent et al., 2021) and in boreal forest (Voříšková et al., 2014; Žifčáková et al., 2017). Our results thus partly confirm the first part of our third hypothesis, that saprotrophs peak in fall or winter and molds+yeast peak in spring (Fig. 7). The ECM fungi appeared to vary more within season, but remained relatively stable across the seasons as hypothesized, whereas, interestingly, the group of other (non-ectomycorrhizal) root-associated fungi followed a trend very similar to saprotrophs across seasons at both sites (Fig. 7). Putative root-associated fungi within Leotiomycetes and Eurotiomycetes (Ascomycota) are often dominant and species-rich in similar tundra systems (Castaño et al., 2023; Clemmensen et al., 2021). While most ECM fungi (Agaricomycetes) lost the genetic capacity to decompose plant cell wall derived OM when evolving a symbiotic lifestyle from saprotrophic ancestors (Miyauchi et al., 2020), the root-associated Ascomycota maintained such capacities (Martino et al., 2018). These fungi likely maintained a versatile life-style, potentially allowing them to shift between being mycorrhizal symbionts during periods with active plant hosts and free-living saprotrophic decomposers when their hosts are inactive (Perotto et al., 2018). The close synchrony with the saprotrophic guild suggests that root-associated fungi respond to environmental conditions in a similar manner, potentially because they shift to a saprotrophic lifestyle during the cold-season, e.g. as decomposers of recently dead ericaceous hair roots, or they maintain their root-association and remain a sink for plant carbon and mineralized N during the cold season.

In contrast to fungal guilds, the abundances of most inorganic N cycling groups decreased towards winter (or fall) at both sites, despite overall higher bacterial abundances in winter compared to summer (Fig. 2, Fig. 7). Overall, several inorganic N cycling genes were significantly negatively related with root-associated fungi or saprotrophs (Fig. 3), matching the opposing seasonal trends. Inorganic N cycling communities showed instead a summer predominance. This pattern was consistent with the seasonal occurrence of archaeal nitrifiers (*amoA*), which predominantly occurred during the spring and summer months (Fig. S7), as well as with the taxonomic seasonal summer ASV affiliated with Nitrososphaerales (Fig. S6, Table S6), a well-known order of ammonia-oxidizers. Together, these results support the second part of our third hypothesis, that inorganic N cycling communities were less prevalent in winter, whereas they likely played a more important role during the summer. Nevertheless, we did not detect a clear succession from nitrifiers (*amoA*) in spring to denitrifiers (*nirK/S* and *nosZI+II*) and nitrate ammonifiers (*nrfA*) in early summer. This could be related to the potentially shorter (day to week scale) temporal dynamics of these functional groups, which were not captured by our monthly sampling. A winter-summer comparison of functional genes based on metagenomic sequencing in an alpine tundra soil revealed similar patterns, with higher relative summer abundances for archaeal ammonia-oxidizers, and most denitrifier and N_2_ fixation genes (Broadbent et al., 2021). However, the summer:winter ratio of *nifH* and *nrfA* genes were zero or near zero in that study. Increasing inorganic N limitation during summer might have led to peaks in N_2_ fixers (*nifH)* and nitrate ammonifiers (*nrfA)* (Fig. 7) for improving N input and N retention, respectively. In line with the peak in *nifH* abundance, taxonomic seasonal summer ASVs were affiliated with known N_2_ fixation taxa (e.g., Rhizobiales and Frankiales) (Fig. S6, Table S6). Similarly, Schostag et al. (2015) reported higher summer abundances of 16S rRNA transcripts for N_2_ fixing microorganisms in Arctic tundra soils from Svalbard.

Over the course of the cold season, functional N gene ratios suggest a transition from higher potential N loss via N_2_O (*nir*:*nosZ* ratio) in early winter to higher N retention potential (*nrfA*:*nir ratio)* in late winter (Fig. 7), potentially linked to the decreasing availability of inorganic N (Fig. 1). The elevated early winter *nir*:*nosZ* ratio coincided with frequent freeze-thaw cycles (Fig. S2), which can increase the availability of easy-degradable OM and nutrients such as nitrate and could therefore enhance N_2_O emissions (Risk et al., 2013; Wertz et al., 2013). With ongoing climate warming, in particular the fall to early winter period warming five times faster in the Arctic than the global average (Rantanen et al., 2022), and more frequent freeze-thaw cycles (Migała et al., 2026), this may further amplify N_2_O emissions. The effect might be even more pronounced as winter communities can exhibit higher activities, higher growth rates and a higher temperature sensitivity compared to summer communities, as shown for respiration and methane production (Lipson et al., 2002; Monson et al., 2006; Poppeliers et al., 2022). While inorganic N, specifically nitrate, may be more available in early winter due to nutrient release from freeze-thaw-induced cell lysis, inorganic N availability declines in late winter (Fig. 1), potentially leading to increasing N limitation. This likely contributes to the observed functional shift towards nitrate ammonifiers (*nrfA*), which can outcompete denitrifiers under nitrate-limiting conditions and may represent an adaptation to conserve N in typically N-limited tundra soils by minimizing the loss via gaseous emissions or leaching (Pandey et al., 2020). A global analysis of *nrfA*:*nir* ratio across terrestrial biomes supports this assumption, with the highest values reported for tundra soils (Saghaï et al., 2023).

### Winter microbial and func+onal trajectories vary under different snow regimes

Winter community dynamics showed broadly similar patterns at both sampling sites, but their timing and magnitudes differed. In Abisko, the sharp fungal decline followed by growth of all three microbial groups occurred in December, whereas in Vassijaure it occurred later, in February, and was less pronounced (Fig. 2). The timing matched the peaks in community turnover that occurred from December to January (Abisko) and from January to February (Vassijaure) (Fig. 5). These site-specific differences were likely driven by contrasting early-winter soil conditions as discussed above.

The difference in timing in winter taxonomic microbial group dynamics also affected the timing of the functional groups (Fig. 7). While in Abisko the potential for N loss via N_2_O (*nir:nosZ* ratio) peaked in fall and early winter and declined in late winter, in Vassijaure, it was also relatively high in late winter. Combined with the lower N retention potential (*nrfA:nir* ratio), the later decline in fungal guilds (February) and the tendency towards a lower potential for N mineralization relative to uptake (SapMY:ECM ratio), Vassijaure may have experienced reduced inorganic N availability in the following spring.

In addition to differences in winter timing, there were overall differences in taxonomic and functional group abundances between the two sites. Although bacterial abundances at both sampling sites showed a similar range, both fungal and archaeal abundances were higher in Vassijaure (Fig. 2). The difference is likely linked to the lower soil pH in Vassijaure. All fungal guilds and N_2_ fixers (*nifH*) were also more abundant in Vassijaure (Fig. 7). In contrast, nitrate ammonifiers (*nrfA*) were considerably more abundant in Abisko, and denitrifiers (*nirS)* and archaeal ammonia-oxidizers (*amoA*) were also detected more frequently there (Fig. S7). The prevalence of nitrate ammonifiers in Abisko might be connected to the pH optimum > 7.5 of the NrfA enzyme (Kajie & Anraku, 1986; Kim et al., 2017), indicating a preference for the higher pH soils at this site. In addition, the slightly lower nitrate content in Abisko might favor nitrate ammonifiers over denitrifiers (Saghaï et al., 2023). The Abisko site exhibited a greater potential for inorganic N cycling overall and the dominance of nitrate ammonifiers suggests a greater potential for N retention, reflected by the higher *nrfA:nir* and lower *nir:nosZ* ratios (Fig. 7). In comparison, Vassijaure appears to be more dominated by ON cycling and mineralization, particularly during the cold season and likely promoted by the more stable conditions due to the higher snow cover as reflected by the positive correlation between snow depth and bacterial and fungal abundances as well as the SapMY:ECM ratio. This finding matches with higher gross N mineralization and nutrient concentrations in Arctic tundra soils under deeper snow (Buckeridge & Grogan, 2010; Xu et al., 2021). In addition, cold season N loss potential (*nir:nosZ* ratio) is higher in Vassijaure. Similarly, in Svalbard heath tundra late winter denitrification potential was higher with deeper snow, while the nitrous oxide reductase potential was not affected by snow depth (Xu et al., 2021). However, the potential N loss during winter at Vassijaure may be partly compensated by a higher potential for summer N input via N_2_ fixation.

### Technical considera+ons

The estimation of microbial community turnover based on DNA analysis might be affected by a higher preservation of DNA in winter due to lower microbial activities, which are caused by lower temperatures and a limited availability of liquid water (Tilston et al., 2010). For rRNA, the half-life considerably increased with decreasing soil temperatures from around 16 days at 4°C to 215 days at −4°C (Schostag et al., 2020). However, our study shows increasing absolute gene abundances over winter for bacteria and fungi (Fig. 2) indicating growth and more importantly newly appearing taxa with a maximum in winter (Fig. 5). Further, community turnover in winter is also supported by the distance-based MEM analysis, which indicates a shift in communities in winter (Fig. S5, Table S5). Thus, we believe that our findings of dynamic microbial communities in winter are well-supported.

## Conclusion

Our study demonstrates that microbial communities in Arctic tundra soils are far more dynamic during winter than previously assumed. All major taxonomic groups (bacteria, fungi, and archaea) underwent shifts in abundance and composition, but these responses were mostly asynchronous. A considerable drop in fungal abundances, likely triggered by repeated freeze-thaw cycles, followed by changes in soil pH, led to a restructuring of winter communities. The timing and magnitude of these shifts differed between sites due to variations in snow cover and soil thermal regimes.

Saprotrophs, molds+yeasts, and other fungal guilds increased in winter or the shoulder seasons, indicating enhanced potential for OM degradation and organic N cycling during the cold period. In contrast, inorganic N cycling groups declined toward winter, despite elevated winter bacterial abundances. By late winter, differences in snow-mediated winter buffering appeared to shift tundra soils between strategies favoring N retention (Abisko) and sustained winter mineralization coupled with higher N loss potential (Vassijaure) with potential legacy effects for the subsequent spring and summer.

Together, our findings underscore that winter is not a biologically dormant season but a critical period that shapes the structure and function of Arctic tundra microbial communities, with significant implications for N turnover potential and winter climate feedbacks.

## Acknowledgements

The authors acknowledge the financial support from the Knut and Alice Wallenberg Foundation (DNR KAW 2020.0126), the Swedish Research Council VR (DNR 2018-04004), and the Kempe Foundation (DNR JCK-1822). The authors also acknowledge support by the National Genomics Infrastructure (NGI) Sweden and Science for Life Laboratory; Illumina sequencing was performed by the SNP&SEQ Technology Platform in Uppsala, and Pacific Biosciences sequencing was performed at Uppsala Genome Center. The SNP&SEQ Platform and Uppsala Genome Center are supported by the Swedish Research Council VR and the Knut and Alice Wallenberg Foundation, Sweden.

We would like to thank Josefine Walz for contribution to the experimental setup and sample collection, Chenxin Feng for help with soil pH measurements, and Magdalena Grudzinska-Sterno for help with the qPCRs. For their help with soil sampling and sample processing, we would like to thank interns and field assistants, including Mattias Dalkvist, Lukas Guth, Mylène Aerts, Miriam, Sebastian, Amber, Sarah H., Maja, Majke, Line, Christoffer Carlborg, Cecilia Thunborg, Sieglinde Kundisch, Max Aaronsson, Lea Traiser, Livia Schulte, Karl Felix Kuhlman, Renée Lejeune, and Louis Schramme.

## Data and code availability

All sequencing data and R code supporting this study will be made publicly available upon publication of the peer-reviewed article.

