## Supplementary Material for "Dynamic winter microbial communities shape nitrogen cycling potential in Arctic tundra soils"

**Table S1:** Primers and conditions for quantification of different microbial taxonomic and functional groups via qPCR.

| Target gene | Primers | Primer conc. (μM) | Sequence | Template DNA (ng)[a] | Cycling conditions[b] | Efficiency | Reference |
|---|---|---|---|---|---|---|---|
| Bacterial 16S rRNA | 515F<br>926R | 1 | GTG YCA GCM GCC GCG GTA A<br>CCG YCA ATT YMT TTR AGT TT | 4 | 95°C 5 min;<br>(95°C 15 s, 50°C 30 s, 72°C 30 s) 35 cycles | 92-97% | (Parada et al., 2016)<br>(Quince et al., 2011) |
| Archaeal 16S rRNA | S-D-Arch-0349-a-S-17<br>S-D-Bact-0785-a-A-21 | 1 | GYG CAS CAG KCG MGA AW<br>GAC TAC HVG GGT ATC TAA TCC | 10 | 95°C 5 min;<br>(95°C 15 s, 56°C 30 s, 72°C 40 s, 80°C 10 s) 40 cycles | 77-78% | (Takai & Horikoshi, 2000)<br>(Herlemann et al., 2011) |
| Fungal ITS | gITS7<br>ITS4/ITS4A | 0.5<br>0.3/0.15 | GTG ART CAT CGA RTC TTT G<br>TCC TCC GCT TAT TGA TAT GC/<br>TCC TCG CCT TAT TGA TAT GC | 4 | 95°C 5 min;<br>(95°C 15 s, 56°C 30 s, 72°C 40 s, 78°C 5 s) 35 cycles | 93-95% | (Ihrmark et al., 2012)<br>(Sterkenburg et al., 2018; White et al., 1990) |
| *nirK* | nirK876F<br>nirK1040R | 0.5 | ATY GGC GGV CAY GGC GA<br>GCC TCG ATC AGR TTR TGG TT | 5 | 95°C 5 min;<br>(95°C 15 s, 63°C(-1°C/cycle) 30 s, 72°C 30s) 6 cycles<br>(95°C 15 s, 58°C 30 s, 72°C 30 s, 80°C 10 s) 35 cycles | 92-97% | (Henry et al., 2004) |
| *nirS* | cd3aFm<br>R3cdm | 0.8 | AAC GYS AAG GAR ACS GG<br>GAS TTC GGR TGS GTC TTS AYG AA | 10 | 95°C 5 min;<br>(95°C 15 s, 65°C(-1°C/cycle) 30 s, 72°C 30s) 6 cycles<br>(95°C 15 s, 60°C 30 s, 72°C 30 s, 80°C 10 s) 35 cycles | 86-88% | (Kandeler et al., 2006) |
| *nosZI* | 1840F<br>2090R | 0.8 | CGC RAC GGC AAS AAG GTS MSS GT<br>CAK RTG CAK SGC RTG GCA GAA | 5 | 95°C 5 min; | 91-95% | (Henry et al., 2006) |

| | | | | | | | |
|---|---|---|---|---|---|---|---|
| | | | | | (95°C 15 s, 65°C(-1°C/cycle) 30 s, 72°C 30s) 6 cycles<br>(95°C 15 s, 60°C 30 s, 72°C 30 s, 80°C 10 s) 35 cycles | | |
| *nosZII* | nosZ-II_F<br>nosZ-II_R | 2 | CTI GGI CCI YTK CAY AC<br>GCI GAR CAR AAI TCB GTR C | 10 | 95°C 7 min;<br>(95°C 15 s, 54°C 30 s, 72°C 40 s, 80°C 10 s) 40 cycles | 84-85% | (Jones et al., 2013) |
| Archaeal *amoA* | Arch-amoA-104F<br>Arch-amoA-616R | 0.5 | GCA GGA GAC TAY ATH TTC TA<br>GCC ATC CAT CTR TAD GTC CA | 15 | 95°C 5 min;<br>(95°C 15 s, 55°C 30 s, 72°C 40 s, 80°C 10 s) 40 cycles | 83-87% | (Alves et al., 2013) |
| Bacterial *amoA* | amoA-1F<br>amoA-2R' | 1 | GGG GTT TCT ACT GGT GGT<br>CCT CKG SAA AGC CTT CTT C | 15 | 95°C 5 min;<br>(95°C 15 s, 55°C 30 s, 72°C 40 s, 80°C 10 s) 40 cycles | 98% | (Rotthauwe et al., 1997)<br>(Okano et al., 2004) |
| *nrfA* | nrfAF2aw_MOD<br>nrfAR1_MOD | 2 | GSI CAR TGY CAY GTI GAR TA<br>GGC ATR TGR CAR TCI RYR CA | 10 | 95°C 5 min;<br>(95°C 30 s, 56°C 30 s, 72°C 30 s, 80°C 10 s) 40 cycles | 76-78% | (Cannon et al., 2019) |
| *nifH* | Ueda19F<br>R6 | 2 | GCI WTY TAY GGI AAR GGI GG<br>GCC ATC ATY TCI CCI GA | 5 | 95°C 5 min;<br>(95°C 15 s, 52°C 30 s, 72°C 40 s, 80°C 10 s) 40 cycles | 83-87% | (Ueda et al., 1995)<br>(Marusina et al., 2001) |

[a] The total volume of each reaction was 15 µL for the bacterial 16S rRNA gene and fungal ITS assays and 12 µL for all other assays.
[b] One of the replicate runs ended with a melt curve: 95°C 15 s and 60 to 95 °C, 10 s increment 0.5 °C.

**Table S2:** Cycling conditions for PCR. Primers and references can be found in Table S1.

| Target gene | Primer conc. (μM) | Template DNA | Cycling conditions |
|---|---|---|---|
| Bacterial 16S rRNA (PCR 1) | 0.25 | 4 ng | 98°C 3 min;<br>(98°C 15 s, 50°C 30 s, 72°C 40 s)<br>25 cycles;<br>72°C 10 min |
| Archaeal 16S rRNA (PCR1) | 0.25 | 10 ng | 98°C 3 min;<br>(98°C 30 s, 65°C 30 s, 72°C 30 s)<br>30 cycles;<br>72°C 10 min |
| 16S rRNA (PCR 2) | 0.2 | 3 μL (PCR1) | 98°C 3 min;<br>(98°C 30 s, 55°C 30 s, 72°C 45 s)<br>8 cycles;<br>72°C 5 min |
| Fungal ITS | 0.5<br>0.3/0.15 | 5 ng | 95°C 5 min;<br>(95°C 30 s, 57°C 30 s, 72°C 30s,)<br>21-35 cycles;<br>72°C 10 min |

**Table S3**: Effect of sampling site and time point or season for each site on microbial community composition (Euclidean distance) for each microbial group based on permutational multivariate analyses of variance (PERMANOVA). Significance was assessed with 999 permutations and block was included as a stratification factor.

| | Bacteria | | Fungi | | Archaea | |
|---|---|---|---|---|---|---|
| Site (n=2) | F=18.011 p<0.001 *** | | F=13.087 p<0.001 *** | | F=12.892 p<0.001 *** | |
| | Abisko | Vassijaure | Abisko | Vassijaure | Abisko | Vassijaure |
| Time point (n=15) | F=1.2477 p=0.018 * | F=1.2957 p=0.002 ** | F=1.0032 p=0.265 | F=0.9737 p=0.239 | F=1.0753 p=0.034 * | F=1.076 p=0.026 * |
| Season (n=5) | F=1.7306 p=0.01 ** | F=2.0849 p<0.001 *** | F=1.2613 p=0.014 * | F=1.2646 p=0.003 ** | F=1.6189 p=0.002 ** | F=1.4039 p=0.013 * |

**Table S4**: Significant temporal scales (MEMs) from distance-based Moran's eigenvector maps (db-MEM) identified by forward selection of partial redundancy analysis accounting for block. MEMs are illustrated in Fig. S5.

| | Bacteria | | Fungi | | Archaea | |
|---|---|---|---|---|---|---|
| | Abisko | Vassijaure | Abisko | Vassijaure | Abisko | Vassijaure |
| Significant MEMs | MEM1 | MEM1<br>MEM2 | MEM1 | MEM1 | MEM2<br>MEM4 | MEM1 |

**Table S5**: Significant environmental variables identified by forward selection in partial redundancy analysis accounting for block. MEMs are illustrated in Fig. S5.

| | Bacteria | | Fungi | | Archaea | |
|---|---|---|---|---|---|---|
| | Abisko | Vassijaure | Abisko | Vassijaure | Abisko | Vassijaure |
| Final model | pH***<br>MEM1*** | pH***<br>MEM1***<br>Tsoil** | pH***<br>MEM1*** | pH***<br>MEM1** | pH***<br>MEM2***<br>MEM4***<br>Snow depth** | pH***<br>MEM1*** |

**Table S6**: Taxonomic classification for seasonal taxa of all three microbial groups (bacteria, fungi, and archaea). Site corresponds to Abisko (abi) and Vassijaure (vas). Rows with red shading correspond to summer taxa and with blue shading to winter taxa.

| No | Site | Season cluster | Microbial group | Phylum | Class | Order | Family | Genus |
|---|---|---|---|---|---|---|---|---|
| 1 | vas | summer | Bacteria | Actinobacteriota | Actinobacteria | Frankiales | Acidothermaceae | Acidothermus |
| 2 | vas | summer | Bacteria | Actinobacteriota | Actinobacteria | Frankiales | Acidothermaceae | Acidothermus |
| 3 | vas | summer | Bacteria | Actinobacteriota | Actinobacteria | Frankiales | Acidothermaceae | Acidothermus |
| 4 | vas | summer | Bacteria | Actinobacteriota | Actinobacteria | Frankiales | Acidothermaceae | Acidothermus |
| 5 | vas | summer | Bacteria | Actinobacteriota | Actinobacteria | Frankiales | Acidothermaceae | Acidothermus |
| 6 | vas | summer | Bacteria | Actinobacteriota | Actinobacteria | Frankiales | Acidothermaceae | Acidothermus |
| 7 | vas | summer | Bacteria | Methylomirabilota | Methylomirabilia | Rokubacteriales | Unclassified | Unclassified |
| 8 | vas | summer | Bacteria | Planctomycetota | Planctomycetes | Gemmatales | Gemmataceae | Unclassified |
| 9 | vas | summer | Bacteria | Acidobacteriota | Acidobacteriae | Subgroup 2 | Unclassified | Unclassified |
| 10 | vas | summer | Bacteria | Acidobacteriota | Acidobacteriae | Subgroup 2 | Unclassified | Unclassified |
| 11 | vas | summer | Bacteria | Acidobacteriota | Acidobacteriae | Subgroup 2 | Unclassified | Unclassified |
| 12 | vas | summer | Bacteria | Proteobacteria | Gammaproteobacteria | Gammaproteobacteria Incertae Sedis | Unknown Family | Acidibacter |
| 13 | vas | summer | Bacteria | Myxococcota | Myxococcia | Myxococcales | Myxococcaceae | Unclassified |
| 14 | vas | summer | Bacteria | Myxococcota | Myxococcia | Myxococcales | Myxococcaceae | Unclassified |
| 15 | vas | summer | Bacteria | Myxococcota | Myxococcia | Myxococcales | Myxococcaceae | Unclassified |
| 16 | vas | summer | Bacteria | Acidobacteriota | Acidobacteriae | Acidobacteriales | Unclassified | Unclassified |
| 17 | vas | summer | Bacteria | Acidobacteriota | Acidobacteriae | Acidobacteriales | Unclassified | Unclassified |
| 18 | vas | summer | Bacteria | Acidobacteriota | Acidobacteriae | Acidobacteriales | Unclassified | Unclassified |
| 19 | vas | summer | Bacteria | Verrucomicrobiota | Verrucomicrobiae | Chthoniobacterales | Chthoniobacteraceae | Candidatus Udaeobacter |
| 20 | vas | summer | Bacteria | Verrucomicrobiota | Verrucomicrobiae | Chthoniobacterales | Chthoniobacteraceae | Candidatus Udaeobacter |
| 21 | vas | summer | Bacteria | Verrucomicrobiota | Verrucomicrobiae | Chthoniobacterales | Chthoniobacteraceae | Candidatus Udaeobacter |
| 22 | vas | summer | Bacteria | Verrucomicrobiota | Verrucomicrobiae | Chthoniobacterales | Chthoniobacteraceae | Candidatus Udaeobacter |
| 23 | vas | summer | Bacteria | Verrucomicrobiota | Verrucomicrobiae | Chthoniobacterales | Xiphinematobacteraceae | Candidatus Xiphinematobacter |
| 24 | vas | summer | Bacteria | Verrucomicrobiota | Verrucomicrobiae | Chthoniobacterales | Chthoniobacteraceae | Candidatus Udaeobacter |
| 25 | vas | summer | Bacteria | Verrucomicrobiota | Verrucomicrobiae | Chthoniobacterales | Chthoniobacteraceae | Candidatus Udaeobacter |
| 26 | vas | summer | Bacteria | Acidobacteriota | Vicinamibacteria | Subgroup 17 | Unclassified | Unclassified |
| 27 | vas | summer | Bacteria | Acidobacteriota | Vicinamibacteria | Subgroup 17 | Unclassified | Unclassified |
| 28 | vas | summer | Bacteria | Acidobacteriota | Vicinamibacteria | Subgroup 17 | Unclassified | Unclassified |
| 29 | vas | summer | Bacteria | Proteobacteria | Alphaproteobacteria | Rhizobiales | Xanthobacteraceae | Unclassified |
| 30 | vas | summer | Bacteria | Proteobacteria | Alphaproteobacteria | Rhizobiales | Xanthobacteraceae | Rhodoplanes |
| 31 | vas | summer | Bacteria | Proteobacteria | Alphaproteobacteria | Rhizobiales | Xanthobacteraceae | Unclassified |
| 32 | vas | summer | Bacteria | Proteobacteria | Alphaproteobacteria | Rhizobiales | Xanthobacteraceae | Unclassified |
| 33 | vas | summer | Bacteria | Proteobacteria | Alphaproteobacteria | Rhizobiales | Amb-16S-1323 | Unclassified |
| 34 | vas | summer | Bacteria | Proteobacteria | Alphaproteobacteria | Rhizobiales | Xanthobacteraceae | Pseudolabrys |
| 35 | vas | summer | Bacteria | Proteobacteria | Alphaproteobacteria | Rhizobiales | Rhizobiales Incertae Sedis | Bauldia |
| 36 | vas | summer | Bacteria | Acidobacteriota | Acidobacteriae | Acidobacteriales | Unclassified | Unclassified |
| 37 | vas | summer | Bacteria | Acidobacteriota | Acidobacteriae | Acidobacteriales | Unclassified | Unclassified |
| 38 | vas | summer | Bacteria | Bacteroidota | Bacteroidia | Chitinophagales | Chitinophagaceae | Puia |
| 39 | vas | summer | Bacteria | Acidobacteriota | Blastocatellia | Pyrinomonadales | Pyrinomonadaceae | RB41 |
| 40 | vas | summer | Bacteria | Acidobacteriota | Vicinamibacteria | Vicinamibacterales | Unclassified | Unclassified |
| 41 | vas | summer | Bacteria | Acidobacteriota | Acidobacteriae | Bryobacterales | Bryobacteraceae | Bryobacter |
| 42 | vas | summer | Bacteria | Acidobacteriota | Acidobacteriae | Bryobacterales | Bryobacteraceae | Bryobacter |
| 43 | vas | summer | Bacteria | Actinobacteriota | Thermoleophilia | Gaiellales | Unclassified | Unclassified |
| 44 | vas | summer | Bacteria | Actinobacteriota | Actinobacteria | Frankiales | Acidothermaceae | Acidothermus |
| 45 | vas | summer | Bacteria | Actinobacteriota | Acidimicrobiia | IMCC26256 | Unclassified | Unclassified |
| 46 | vas | summer | Bacteria | Actinobacteriota | Thermoleophilia | Solirubrobacterales | Solirubrobacteraceae | Conexibacter |
| 47 | vas | summer | Bacteria | Actinobacteriota | Thermoleophilia | Solirubrobacterales | Solirubrobacteraceae | Conexibacter |

| | | | | | | | | |
|---|---|---|---|---|---|---|---|---|
| 48 | vas | summer | Bacteria | Actinobacteriota | Thermoleophilia | Solirubrobacterales | 67-14 | Unclassified |
| 49 | vas | summer | Bacteria | Actinobacteriota | Thermoleophilia | Gaiellales | Gaiellaceae | Gaiella |
| 50 | vas | summer | Bacteria | Actinobacteriota | Thermoleophilia | Gaiellales | Unclassified | Unclassified |
| 51 | vas | summer | Bacteria | Actinobacteriota | Thermoleophilia | Gaiellales | Unclassified | Unclassified |
| 52 | vas | summer | Bacteria | Actinobacteriota | Thermoleophilia | Solirubrobacterales | 67-14 | Unclassified |
| 53 | vas | summer | Bacteria | Actinobacteriota | Thermoleophilia | Solirubrobacterales | 67-14 | Unclassified |
| 54 | vas | summer | Bacteria | Actinobacteriota | Thermoleophilia | Solirubrobacterales | Solirubrobacteraceae | Solirubrobacter |
| 55 | vas | summer | Bacteria | Actinobacteriota | Acidimicrobiia | IMCC26256 | Unclassified | Unclassified |
| 56 | vas | summer | Bacteria | Actinobacteriota | Thermoleophilia | Solirubrobacterales | Solirubrobacteraceae | Unclassified |
| 57 | vas | summer | Bacteria | Actinobacteriota | Thermoleophilia | Solirubrobacterales | Solirubrobacteraceae | Unclassified |
| 58 | vas | summer | Bacteria | Actinobacteriota | Actinobacteria | Frankiales | Sporichthyaceae | Unclassified |
| 59 | vas | summer | Bacteria | Actinobacteriota | Actinobacteria | Frankiales | Frankiaceae | Jatrophihabitans |
| 60 | vas | summer | Bacteria | Actinobacteriota | Actinobacteria | Frankiales | Frankiaceae | Jatrophihabitans |
| 61 | vas | summer | Bacteria | Actinobacteriota | Acidimicrobiia | Unclassified | Unclassified | Unclassified |
| 62 | vas | summer | Bacteria | Actinobacteriota | Actinobacteria | Frankiales | Sporichthyaceae | Unclassified |
| 63 | vas | summer | Bacteria | Proteobacteria | Gammaproteobacteria | Burkholderiales | A21b | Unclassified |
| 64 | vas | summer | Bacteria | Proteobacteria | Gammaproteobacteria | Burkholderiales | Nitrosomonadaceae | mle1-7 |
| 65 | vas | summer | Bacteria | Proteobacteria | Gammaproteobacteria | Burkholderiales | A21b | Unclassified |
| 66 | vas | summer | Bacteria | Proteobacteria | Gammaproteobacteria | Burkholderiales | SC-I-84 | Unclassified |
| 67 | vas | summer | Bacteria | Proteobacteria | Gammaproteobacteria | Burkholderiales | Nitrosomonadaceae | Ellin6067 |
| 68 | vas | summer | Bacteria | Chloroflexi | TK10 | Unclassified | Unclassified | Unclassified |
| 69 | vas | summer | Bacteria | Chloroflexi | Gitt-GS-136 | Unclassified | Unclassified | Unclassified |
| 1 | vas | winter | Bacteria | Actinobacteriota | Actinobacteria | Frankiales | Acidothermaceae | Acidothermus |
| 2 | vas | winter | Bacteria | Planctomycetota | Phycisphaerae | Tepidisphaerales | WD2101 soil group | Unclassified |
| 3 | vas | winter | Bacteria | Planctomycetota | Phycisphaerae | Tepidisphaerales | WD2101 soil group | Unclassified |
| 4 | vas | winter | Bacteria | Planctomycetota | Phycisphaerae | Tepidisphaerales | WD2101 soil group | Unclassified |
| 5 | vas | winter | Bacteria | Planctomycetota | Planctomycetes | Isosphaerales | Isosphaeraceae | Aquisphaera |
| 6 | vas | winter | Bacteria | Planctomycetota | Planctomycetes | Isosphaerales | Isosphaeraceae | Aquisphaera |
| 7 | vas | winter | Bacteria | Planctomycetota | Planctomycetes | Isosphaerales | Isosphaeraceae | Aquisphaera |
| 8 | vas | winter | Bacteria | Planctomycetota | Planctomycetes | Isosphaerales | Isosphaeraceae | Unclassified |
| 9 | vas | winter | Bacteria | Planctomycetota | Planctomycetes | Isosphaerales | Isosphaeraceae | Unclassified |
| 10 | vas | winter | Bacteria | Planctomycetota | Planctomycetes | Isosphaerales | Isosphaeraceae | Tundrisphaera |
| 11 | vas | winter | Bacteria | Planctomycetota | Planctomycetes | Isosphaerales | Isosphaeraceae | Tundrisphaera |
| 12 | vas | winter | Bacteria | Planctomycetota | Planctomycetes | Isosphaerales | Isosphaeraceae | Tundrisphaera |
| 13 | vas | winter | Bacteria | Planctomycetota | Planctomycetes | Gemmatales | Gemmataceae | Gemmata |
| 14 | vas | winter | Bacteria | Planctomycetota | Planctomycetes | Isosphaerales | Isosphaeraceae | Aquisphaera |
| 15 | vas | winter | Bacteria | Planctomycetota | Planctomycetes | Isosphaerales | Isosphaeraceae | Aquisphaera |
| 16 | vas | winter | Bacteria | Proteobacteria | Alphaproteobacteria | Reyranellales | Reyranellaceae | Reyranella |
| 17 | vas | winter | Bacteria | WPS-2 | Unclassified | Unclassified | Unclassified | Unclassified |
| 18 | vas | winter | Bacteria | Verrucomicrobiota | Verrucomicrobiae | Opitutales | Opitutaceae | Lacunisphaera |
| 19 | vas | winter | Bacteria | Proteobacteria | Gammaproteobacteria | Gammaproteobacteria Incertae Sedis | Unknown Family | Acidibacter |
| 20 | vas | winter | Bacteria | Proteobacteria | Gammaproteobacteria | JG36-TzT-191 | Unclassified | Unclassified |
| 21 | vas | winter | Bacteria | Proteobacteria | Gammaproteobacteria | WD260 | Unclassified | Unclassified |
| 22 | vas | winter | Bacteria | Proteobacteria | Gammaproteobacteria | WD260 | Unclassified | Unclassified |
| 23 | vas | winter | Bacteria | Proteobacteria | Gammaproteobacteria | WD260 | Unclassified | Unclassified |
| 24 | vas | winter | Bacteria | Proteobacteria | Gammaproteobacteria | WD260 | Unclassified | Unclassified |
| 25 | vas | winter | Bacteria | Proteobacteria | Gammaproteobacteria | JG36-TzT-191 | Unclassified | Unclassified |
| 26 | vas | winter | Bacteria | Proteobacteria | Gammaproteobacteria | JG36-TzT-191 | Unclassified | Unclassified |
| 27 | vas | winter | Bacteria | Proteobacteria | Gammaproteobacteria | JG36-TzT-191 | Unclassified | Unclassified |
| 28 | vas | winter | Bacteria | Verrucomicrobiota | Verrucomicrobiae | Chthoniobacterales | Xiphinematobacteraceae | Candidatus Xiphinematobacter |
| 29 | vas | winter | Bacteria | Verrucomicrobiota | Verrucomicrobiae | Chthoniobacterales | Chthoniobacteraceae | Chthoniobacter |
| 30 | vas | winter | Bacteria | Verrucomicrobiota | Verrucomicrobiae | Chthoniobacterales | Chthoniobacteraceae | Unclassified |

| | | | | | | | | |
|---|---|---|---|---|---|---|---|---|
| 31 | vas | winter | Bacteria | Planctomycetota | vadinHA49 | Unclassified | Unclassified | Unclassified |
| 32 | vas | winter | Bacteria | Proteobacteria | Alphaproteobacteria | Elsterales | Unclassified | Unclassified |
| 33 | vas | winter | Bacteria | Proteobacteria | Alphaproteobacteria | Elsterales | Unclassified | Unclassified |
| 34 | vas | winter | Bacteria | Proteobacteria | Alphaproteobacteria | Rhizobiales | Xanthobacteraceae | Unclassified |
| 35 | vas | winter | Bacteria | Proteobacteria | Alphaproteobacteria | Acetobacterales | Acetobacteraceae | Unclassified |
| 36 | vas | winter | Bacteria | Proteobacteria | Alphaproteobacteria | Acetobacterales | Acetobacteraceae | Rhodopila |
| 37 | vas | winter | Bacteria | Proteobacteria | Alphaproteobacteria | Acetobacterales | Acetobacteraceae | Unclassified |
| 38 | vas | winter | Bacteria | Proteobacteria | Alphaproteobacteria | Acetobacterales | Acetobacteraceae | Acidisoma |
| 39 | vas | winter | Bacteria | Proteobacteria | Alphaproteobacteria | Acetobacterales | Acetobacteraceae | Acidicaldus |
| 40 | vas | winter | Bacteria | Proteobacteria | Alphaproteobacteria | Rhizobiales | Beijerinckiaceae | Roseiarcus |
| 41 | vas | winter | Bacteria | Proteobacteria | Alphaproteobacteria | Rhizobiales | Beijerinckiaceae | Roseiarcus |
| 42 | vas | winter | Bacteria | Proteobacteria | Alphaproteobacteria | Micropepsales | Micropepsaceae | Unclassified |
| 43 | vas | winter | Bacteria | Proteobacteria | Gammaproteobacteria | Xanthomonadales | Rhodanobacteraceae | Rhodanobacter |
| 44 | vas | winter | Bacteria | Acidobacteriota | Acidobacteriae | Acidobacteriales | Unclassified | Unclassified |
| 45 | vas | winter | Bacteria | Acidobacteriota | Acidobacteriae | Acidobacteriales | Acidobacteriaceae (Subgroup 1) | Granulicella |
| 46 | vas | winter | Bacteria | Acidobacteriota | Acidobacteriae | Acidobacteriales | Acidobacteriaceae (Subgroup 1) | Granulicella |
| 47 | vas | winter | Bacteria | Acidobacteriota | Acidobacteriae | Acidobacteriales | Acidobacteriaceae (Subgroup 1) | Granulicella |
| 48 | vas | winter | Bacteria | Acidobacteriota | Acidobacteriae | Acidobacteriales | Acidobacteriaceae (Subgroup 1) | Granulicella |
| 49 | vas | winter | Bacteria | Acidobacteriota | Acidobacteriae | Acidobacteriales | Acidobacteriaceae (Subgroup 1) | Granulicella |
| 50 | vas | winter | Bacteria | Verrucomicrobiota | Chlamydiae | Chlamydiales | Simkaniaceae | Unclassified |
| 51 | vas | winter | Bacteria | NB1-j | Unclassified | Unclassified | Unclassified | Unclassified |
| 52 | vas | winter | Bacteria | Acidobacteriota | Subgroup 22 | Unclassified | Unclassified | Unclassified |
| 53 | vas | winter | Bacteria | Bacteroidota | Bacteroidia | Chitinophagales | Chitinophagaceae | Puia |
| 54 | vas | winter | Bacteria | Bacteroidota | Bacteroidia | Chitinophagales | Chitinophagaceae | Puia |
| 55 | vas | winter | Bacteria | Acidobacteriota | Vicinamibacteria | Vicinamibacterales | Unclassified | Unclassified |
| 56 | vas | winter | Bacteria | Armatimonadota | Chthonomonadetes | Chthonomonadales | Chthonomonadaceae | Chthonomonas |
| 57 | vas | winter | Bacteria | Acidobacteriota | Acidobacteriae | Bryobacterales | Bryobacteraceae | Bryobacter |
| 58 | vas | winter | Bacteria | Acidobacteriota | Acidobacteriae | Bryobacterales | Bryobacteraceae | Bryobacter |
| 59 | vas | winter | Bacteria | Acidobacteriota | Acidobacteriae | Solibacterales | Solibacteraceae | Candidatus Solibacter |
| 60 | vas | winter | Bacteria | Acidobacteriota | Acidobacteriae | Solibacterales | Solibacteraceae | Candidatus Solibacter |
| 61 | vas | winter | Bacteria | Acidobacteriota | Acidobacteriae | Solibacterales | Solibacteraceae | Candidatus Solibacter |
| 62 | vas | winter | Bacteria | Acidobacteriota | Acidobacteriae | Solibacterales | Solibacteraceae | Candidatus Solibacter |
| 63 | vas | winter | Bacteria | Acidobacteriota | Acidobacteriae | Solibacterales | Solibacteraceae | Candidatus Solibacter |
| 64 | vas | winter | Bacteria | Acidobacteriota | Acidobacteriae | Solibacterales | Solibacteraceae | Candidatus Solibacter |
| 65 | vas | winter | Bacteria | Actinobacteriota | Actinobacteria | Frankiales | Acidothermaceae | Acidothermus |
| 66 | vas | winter | Bacteria | Acidobacteriota | Acidobacteriae | Subgroup 2 | Unclassified | Unclassified |
| 67 | vas | winter | Bacteria | Proteobacteria | Gammaproteobacteria | Burkholderiales | Comamonadaceae | Piscinibacter |
| 68 | vas | winter | Bacteria | Proteobacteria | Gammaproteobacteria | Burkholderiales | Comamonadaceae | Piscinibacter |
| 69 | vas | winter | Bacteria | Bacteroidota | Bacteroidia | Cytophagales | Microscillaceae | OLB12 |
| 1 | abi | summer | Bacteria | Actinobacteriota | Actinobacteria | Micromonosporales | Micromonosporaceae | Actinoplanes |
| 2 | abi | summer | Bacteria | Actinobacteriota | Actinobacteria | Micromonosporales | Micromonosporaceae | Luedemannella |
| 3 | abi | summer | Bacteria | Actinobacteriota | MB-A2-108 | Unclassified | Unclassified | Unclassified |
| 4 | abi | summer | Bacteria | Myxococcota | Myxococcia | Myxococcales | Myxococcaceae | Unclassified |
| 5 | abi | summer | Bacteria | Myxococcota | Myxococcia | Myxococcales | Myxococcaceae | Unclassified |
| 6 | abi | summer | Bacteria | Acidobacteriota | Acidobacteriae | Acidobacteriales | Unclassified | Unclassified |
| 7 | abi | summer | Bacteria | Proteobacteria | Gammaproteobacteria | Gammaproteobacteria Incertae Sedis | Unknown Family | Acidibacter |
| 8 | abi | summer | Bacteria | Acidobacteriota | Acidobacteriae | Acidobacteriales | Unclassified | Unclassified |
| 9 | abi | summer | Bacteria | Planctomycetota | Phycisphaerae | Tepidisphaerales | WD2101 soil group | Unclassified |
| 10 | abi | summer | Bacteria | Verrucomicrobiota | Verrucomicrobiae | Chthoniobacterales | Chthoniobacteraceae | Chthoniobacter |
| 11 | abi | summer | Bacteria | Acidobacteriota | Vicinamibacteria | Subgroup 17 | Unclassified | Unclassified |
| 12 | abi | summer | Bacteria | Acidobacteriota | Vicinamibacteria | Subgroup 17 | Unclassified | Unclassified |
| 13 | abi | summer | Bacteria | Acidobacteriota | Vicinamibacteria | Subgroup 17 | Unclassified | Unclassified |

| | | | | | | | | |
|---|---|---|---|---|---|---|---|---|
| 14 | abi | summer | Bacteria | Acidobacteriota | Vicinamibacteria | Vicinamibacterales | Unclassified | Unclassified |
| 15 | abi | summer | Bacteria | Planctomycetota | Planctomycetes | Pirellulales | Pirellulaceae | Pir4 lineage |
| 16 | abi | summer | Bacteria | Proteobacteria | Alphaproteobacteria | Rhizobiales | Hyphomicrobiaceae | Hyphomicrobium |
| 17 | abi | summer | Bacteria | Proteobacteria | Alphaproteobacteria | Rhizobiales | Unclassified | Unclassified |
| 18 | abi | summer | Bacteria | Proteobacteria | Alphaproteobacteria | Rhizobiales | Xanthobacteraceae | Rhodoplanes |
| 19 | abi | summer | Bacteria | Proteobacteria | Alphaproteobacteria | Rhizobiales | Xanthobacteraceae | Unclassified |
| 20 | abi | summer | Bacteria | Proteobacteria | Alphaproteobacteria | Rhizobiales | Xanthobacteraceae | Bradyrhizobium |
| 21 | abi | summer | Bacteria | Proteobacteria | Alphaproteobacteria | Rhizobiales | Rhizobiales Incertae Sedis | Unclassified |
| 22 | abi | summer | Bacteria | Acidobacteriota | Acidobacteriae | Acidobacteriales | Unclassified | Unclassified |
| 23 | abi | summer | Bacteria | Acidobacteriota | Acidobacteriae | Acidobacteriales | Unclassified | Unclassified |
| 24 | abi | summer | Bacteria | Bacteroidota | Bacteroidia | Chitinophagales | Chitinophagaceae | Terrimonas |
| 25 | abi | summer | Bacteria | Desulfobacterota | Unclassified | Unclassified | Unclassified | Unclassified |
| 26 | abi | summer | Bacteria | Acidobacteriota | Blastocatellia | Pyrinomonadales | Pyrinomonadaceae | RB41 |
| 27 | abi | summer | Bacteria | Acidobacteriota | Blastocatellia | Pyrinomonadales | Pyrinomonadaceae | RB41 |
| 28 | abi | summer | Bacteria | Acidobacteriota | Blastocatellia | Pyrinomonadales | Pyrinomonadaceae | RB41 |
| 29 | abi | summer | Bacteria | Acidobacteriota | Blastocatellia | Pyrinomonadales | Pyrinomonadaceae | RB41 |
| 30 | abi | summer | Bacteria | Acidobacteriota | Vicinamibacteria | Vicinamibacterales | Unclassified | Unclassified |
| 31 | abi | summer | Bacteria | Acidobacteriota | Vicinamibacteria | Vicinamibacterales | Unclassified | Unclassified |
| 32 | abi | summer | Bacteria | Actinobacteriota | Acidimicrobiia | Microtrichales | Ilumatobacteraceae | CL500-29 marine group |
| 33 | abi | summer | Bacteria | Actinobacteriota | Acidimicrobiia | IMCC26256 | Unclassified | Unclassified |
| 34 | abi | summer | Bacteria | Actinobacteriota | Acidimicrobiia | IMCC26256 | Unclassified | Unclassified |
| 35 | abi | summer | Bacteria | Actinobacteriota | Thermoleophilia | Solirubrobacterales | Solirubrobacteraceae | Conexibacter |
| 36 | abi | summer | Bacteria | Actinobacteriota | Thermoleophilia | Solirubrobacterales | 67-14 | Unclassified |
| 37 | abi | summer | Bacteria | Actinobacteriota | Thermoleophilia | Gaiellales | Gaiellaceae | Gaiella |
| 38 | abi | summer | Bacteria | Actinobacteriota | Thermoleophilia | Gaiellales | Unclassified | Unclassified |
| 39 | abi | summer | Bacteria | Actinobacteriota | Thermoleophilia | Solirubrobacterales | Solirubrobacteraceae | Solirubrobacter |
| 40 | abi | summer | Bacteria | Actinobacteriota | Actinobacteria | Propionibacteriales | Nocardioidaceae | Nocardioides |
| 41 | abi | summer | Bacteria | Actinobacteriota | Actinobacteria | Frankiales | Sporichthyaceae | Unclassified |
| 42 | abi | summer | Bacteria | Actinobacteriota | Actinobacteria | Kineosporiales | Kineosporiaceae | Kineosporia |
| 43 | abi | summer | Bacteria | Actinobacteriota | Actinobacteria | Frankiales | Nakamurellaceae | Nakamurella |
| 44 | abi | summer | Bacteria | Proteobacteria | Gammaproteobacteria | Burkholderiales | SC-I-84 | Unclassified |
| 45 | abi | summer | Bacteria | Proteobacteria | Gammaproteobacteria | Burkholderiales | Nitrosomonadaceae | GOUTA6 |
| 46 | abi | summer | Bacteria | Proteobacteria | Gammaproteobacteria | Burkholderiales | Nitrosomonadaceae | Ellin6067 |
| 47 | abi | summer | Bacteria | Proteobacteria | Gammaproteobacteria | Burkholderiales | Nitrosomonadaceae | Ellin6067 |
| 48 | abi | summer | Bacteria | Chloroflexi | TK10 | Unclassified | Unclassified | Unclassified |
| 49 | abi | summer | Bacteria | Actinobacteriota | Actinobacteria | Corynebacteriales | Mycobacteriaceae | Mycobacterium |
| 50 | abi | summer | Bacteria | Actinobacteriota | Actinobacteria | Pseudonocardiales | Pseudonocardiaceae | Pseudonocardia |
| 51 | abi | summer | Bacteria | Firmicutes | Bacilli | Bacillales | Planococcaceae | Psychrobacillus |
| 1 | abi | winter | Bacteria | Verrucomicrobiota | Verrucomicrobiae | Methylacidiphilales | Methylacidiphilaceae | Unclassified |
| 2 | abi | winter | Bacteria | Planctomycetota | Planctomycetes | Gemmatales | Gemmataceae | Gemmata |
| 3 | abi | winter | Bacteria | Planctomycetota | Planctomycetes | Isosphaerales | Isosphaeraceae | Aquisphaera |
| 4 | abi | winter | Bacteria | Proteobacteria | Alphaproteobacteria | Reyranellales | Reyranellaceae | Reyranella |
| 5 | abi | winter | Bacteria | Verrucomicrobiota | Verrucomicrobiae | Opitutales | Opitutaceae | Lacunisphaera |
| 6 | abi | winter | Bacteria | Proteobacteria | Gammaproteobacteria | Gammaproteobacteria Incertae Sedis | Unknown Family | Acidibacter |
| 7 | abi | winter | Bacteria | Proteobacteria | Gammaproteobacteria | Gammaproteobacteria Incertae Sedis | Unknown Family | Acidibacter |
| 8 | abi | winter | Bacteria | Proteobacteria | Gammaproteobacteria | R7C24 | Unclassified | Unclassified |
| 9 | abi | winter | Bacteria | Proteobacteria | Gammaproteobacteria | Gammaproteobacteria Incertae Sedis | Unknown Family | Acidibacter |
| 10 | abi | winter | Bacteria | Proteobacteria | Alphaproteobacteria | Acetobacterales | Acetobacteraceae | Acidisoma |
| 11 | abi | winter | Bacteria | Proteobacteria | Alphaproteobacteria | Rhizobiales | KF-JG30-B3 | Unclassified |
| 12 | abi | winter | Bacteria | Proteobacteria | Gammaproteobacteria | Xanthomonadales | Rhodanobacteraceae | Rhodanobacter |
| 13 | abi | winter | Bacteria | Acidobacteriota | Acidobacteriae | Acidobacteriales | Acidobacteriaceae (Subgroup 1) | Granulicella |
| 14 | abi | winter | Bacteria | Bacteroidota | Bacteroidia | Chitinophagales | Chitinophagaceae | Unclassified |

| | | | | | | | | |
|---|---|---|---|---|---|---|---|---|
| 15 | abi | winter | Bacteria | Bacteroidota | Bacteroidia | Chitinophagales | Chitinophagaceae | Unclassified |
| 16 | abi | winter | Bacteria | Bacteroidota | Bacteroidia | Chitinophagales | Chitinophagaceae | Ferruginibacter |
| 17 | abi | winter | Bacteria | Acidobacteriota | Blastocatellia | Pyrinomonadales | Pyrinomonadaceae | RB41 |
| 18 | abi | winter | Bacteria | Armatimonadota | Chthonomonadetes | Chthonomonadales | Chthonomonadaceae | Chthonomonas |
| 19 | abi | winter | Bacteria | Acidobacteriota | Acidobacteriae | Bryobacterales | Bryobacteraceae | Bryobacter |
| 20 | abi | winter | Bacteria | Actinobacteriota | Thermoleophilia | Gaiellales | Unclassified | Unclassified |
| 21 | abi | winter | Bacteria | Proteobacteria | Gammaproteobacteria | Burkholderiales | Comamonadaceae | Leptothrix |
| 22 | abi | winter | Bacteria | Actinobacteriota | Actinobacteria | Pseudonocardiales | Pseudonocardiaceae | Pseudonocardia |
| 23 | abi | winter | Bacteria | Bacteroidota | Bacteroidia | Cytophagales | Microscillaceae | OLB12 |
| 1 | abi | summer | k__Fungi | p__Basidiomycota | c__Agaricomycetes | o__Russulales | f__Russulaceae | g__Russula |
| 1 | abi | winter | k__Fungi | p__Ascomycota | c__Eurotiomycetes | o__Chaetothyriales | f__Herpotrichiellaceae | g__Herpotrichiellaceae_gen_unspecified |
| 1 | vas | winter | k__Fungi | p__Ascomycota | c__Leotiomycetes | o__Helotiales | f__Hyaloscyphaceae | g__Hyaloscypha |
| 1 | vas | summer | k__Fungi | p__Basidiomycota | c__Agaricomycetes | o__Russulales | f__Russulaceae | g__Russula |
| 2 | vas | summer | k__Fungi | p__Ascomycota | c__Sordariomycetes | o__Sordariomycetes_ord_unspecified | f__Sordariomycetes_fam_unspecified | g__Sordariomycetes_gen_unspecified |
| 1 | abi | summer | Archaea | Crenarchaeota | Nitrososphaeria | Nitrososphaerales | Nitrososphaeraceae | Candidatus Nitrososphaera |
| 2 | abi | summer | Archaea | Nanoarchaeota | Nanoarchaeia | Woesearchaeales | Unclassified | Unclassified |
| 3 | abi | summer | Archaea | Crenarchaeota | Nitrososphaeria | Nitrososphaerales | Nitrososphaeraceae | Unclassified |
| 4 | abi | summer | Archaea | Crenarchaeota | Nitrososphaeria | Nitrososphaerales | Nitrososphaeraceae | Unclassified |
| 5 | abi | summer | Archaea | Nanoarchaeota | Nanoarchaeia | Woesearchaeales | Unclassified | Unclassified |
| 1 | abi | winter | Archaea | Nanoarchaeota | Nanoarchaeia | Woesearchaeales | Unclassified | Unclassified |
| 2 | abi | winter | Archaea | Nanoarchaeota | Nanoarchaeia | Woesearchaeales | Unclassified | Unclassified |
| 3 | abi | winter | Archaea | Crenarchaeota | Nitrososphaeria | Group 1.1c | Unclassified | Unclassified |
| 4 | abi | winter | Archaea | Micrarchaeota | Micrarchaeia | Micrarchaeales | Unclassified | Unclassified |
| 5 | abi | winter | Archaea | Nanoarchaeota | Nanoarchaeia | Woesearchaeales | Unclassified | Unclassified |
| 1 | vas | summer | Archaea | Thermoplasmatota | Thermoplasmata | Marine Group II | Unclassified | Unclassified |
| 2 | vas | summer | Archaea | Thermoplasmatota | Thermoplasmata | Marine Group II | Unclassified | Unclassified |
| 3 | vas | summer | Archaea | Crenarchaeota | Nitrososphaeria | Group 1.1c | Unclassified | Unclassified |
| 4 | vas | summer | Archaea | Crenarchaeota | Nitrososphaeria | Group 1.1c | Unclassified | Unclassified |
| 1 | vas | winter | Archaea | Nanoarchaeota | Nanoarchaeia | Woesearchaeales | Unclassified | Unclassified |
| 2 | vas | winter | Archaea | Nanoarchaeota | Nanoarchaeia | Woesearchaeales | Unclassified | Unclassified |
| 3 | vas | winter | Archaea | Nanoarchaeota | Nanoarchaeia | Woesearchaeales | Unclassified | Unclassified |
| 4 | vas | winter | Archaea | Crenarchaeota | Nitrososphaeria | Group 1.1c | Unclassified | Unclassified |
| 5 | vas | winter | Archaea | Crenarchaeota | Nitrososphaeria | Group 1.1c | Unclassified | Unclassified |

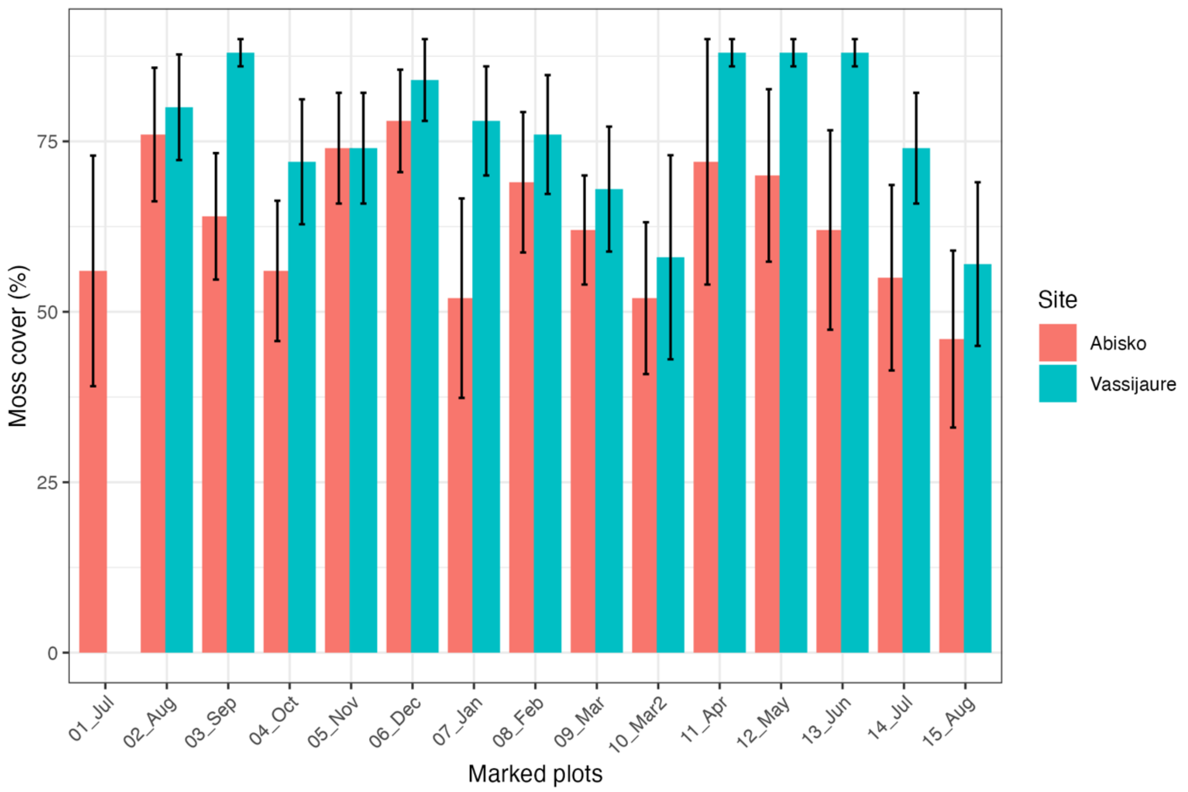


**Figure S1**: Moss cover (%) in marked plots for the one-year sampling campaign at two tundra sites in Abisko and Vassijaure. Note that for all plots, moss cover was measured at the same time in summer 2019 before the actual start of the soil sampling. Values for Vassijaure for time point "01_Jul" plots are not available. Moss cover differed significantly between the two sites (one-way ANOVA, $p<0.001$).

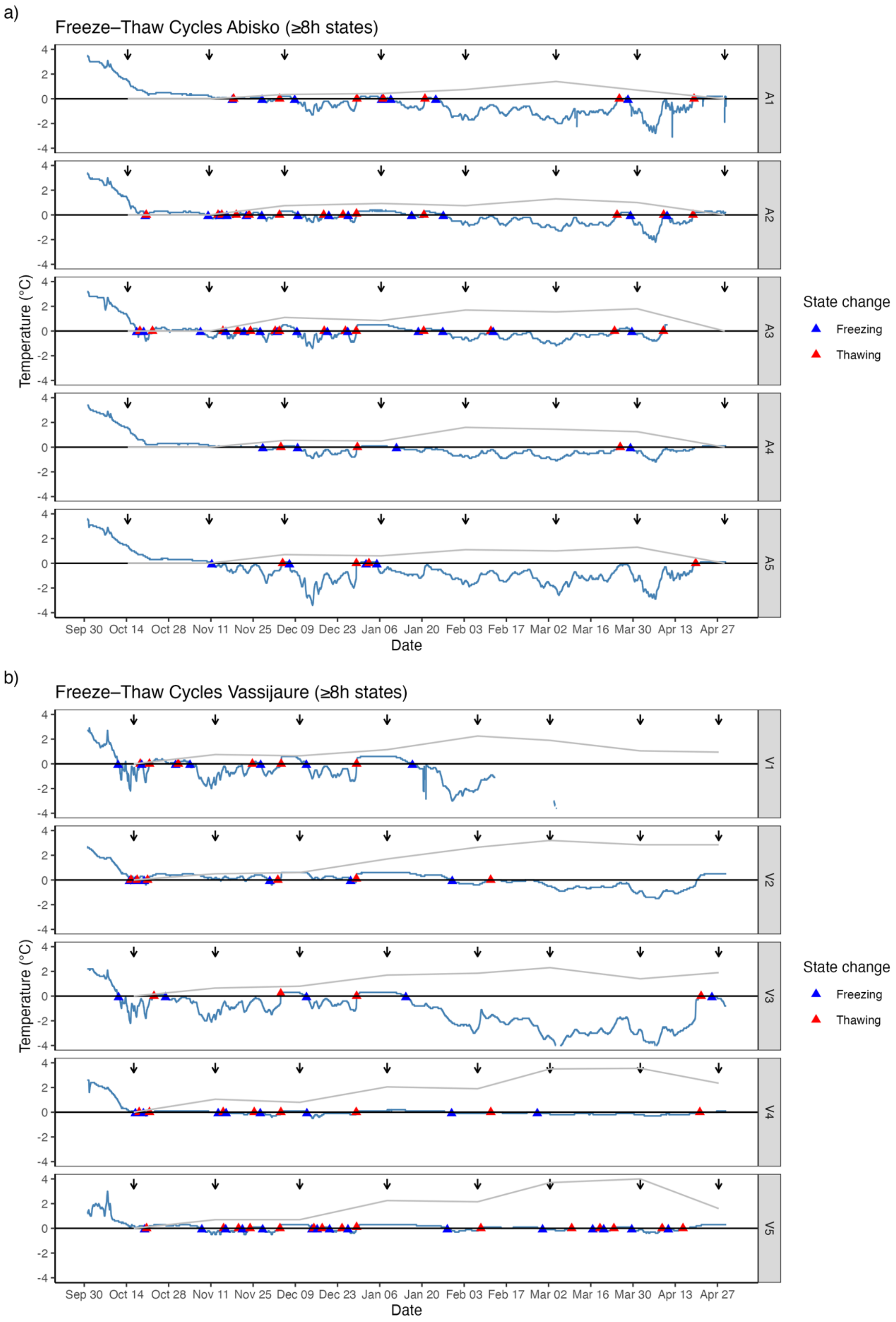


**Figure S2:** Soil temperature profiles (blue line) from October 2019 to April 2020 at both tundra sites for each block at 5 cm depth. Temperature transitions are marked with blue (freezing) and red (thawing) triangles. The arrows indicate the timepoints of the soi sampling for microbial community analysis. The snow depth in the respective sampling plot is shown as grey line.

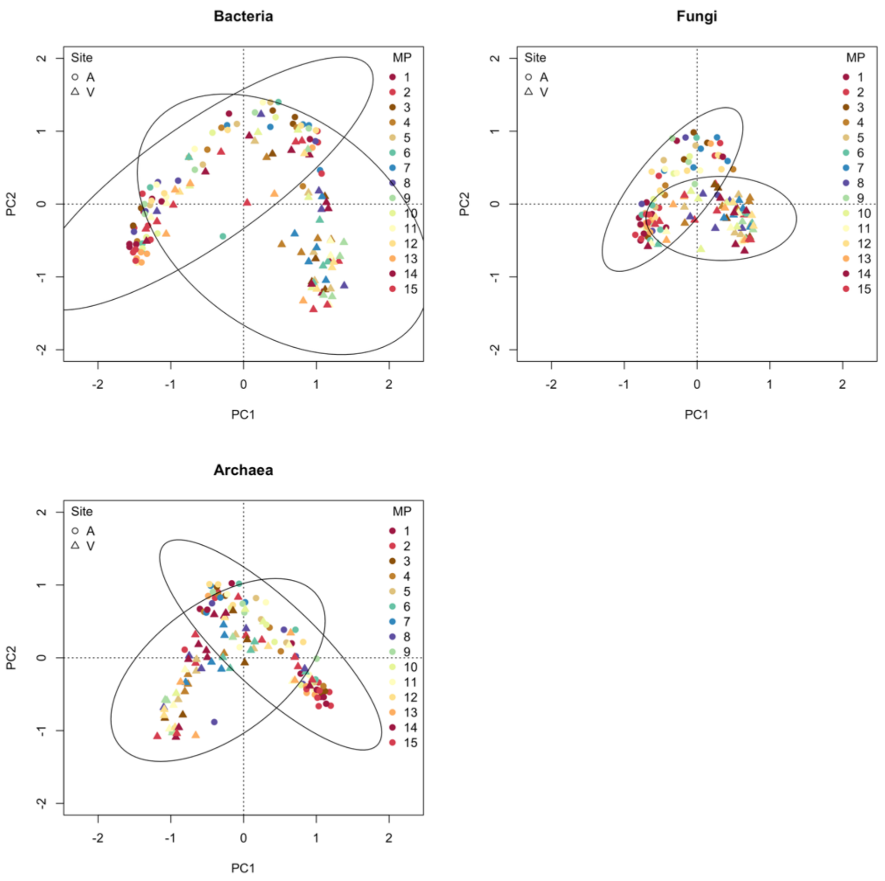


**Figure S3**: Principal component analysis (PCA) of microbial communities including both tundra sites. Symbol shapes denote site (A – Abisko, V – Vassijaure), symbol colors denote time point (MP), and ellipses indicate within-site dispersion in PCA space, scaled to encompass approximately 95% of the variability.

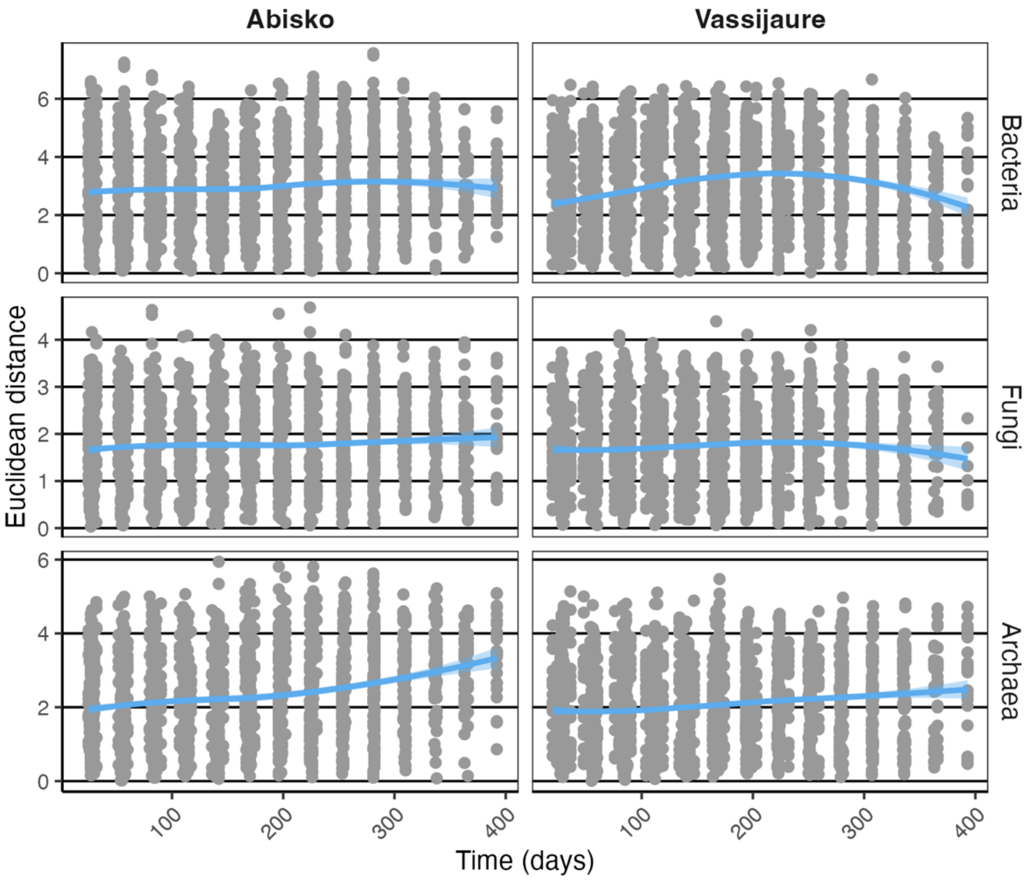


**Figure S4**: Time-decay relationship between time intervalls and microbial community dissimilarity for bacterial, fungal, and archaeal communities based on clr-transformed relative ASV/species abundances during one year from July 2019 to August 2020 at two tundra sites in Abisko and Vassijaure. Pairwise Euclidean distances were plotted against the time differences between sampling time points. The blue line indicates a locally weighted regression (LOESS) smooth, and the blue-shaded area reflects the 95% confidence intervals.

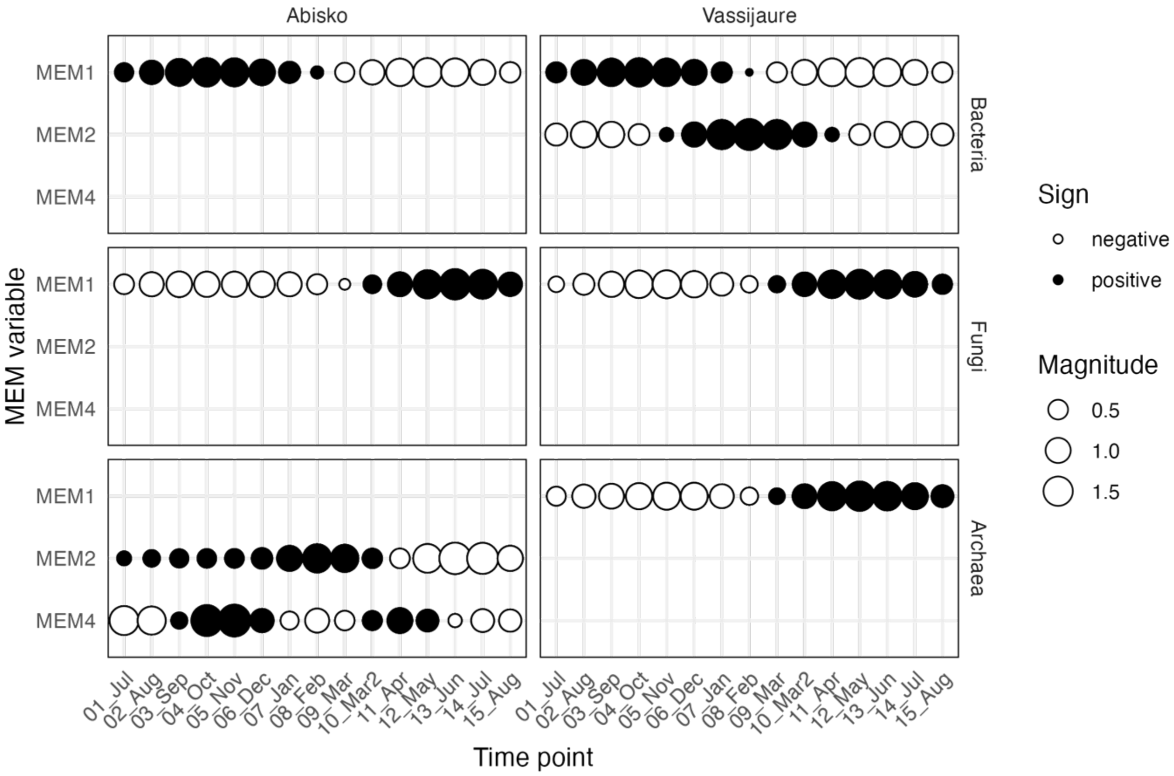


**Figure S5**: Temporal pattern of significant Moran's eigenvector maps (MEM variables) identified by forward selection of partial redundancy analysis of microbial community compositions accounting for block. Points show time points and point size and color denote lower (negative) and higher (positive) eigenvector values. MEM1 represents broad-scale temporal variation with a community shift in winter, while MEM2 and MEM4 represent intermediate temporal variations with MEM4 showing variation similar to seasonal shifts.

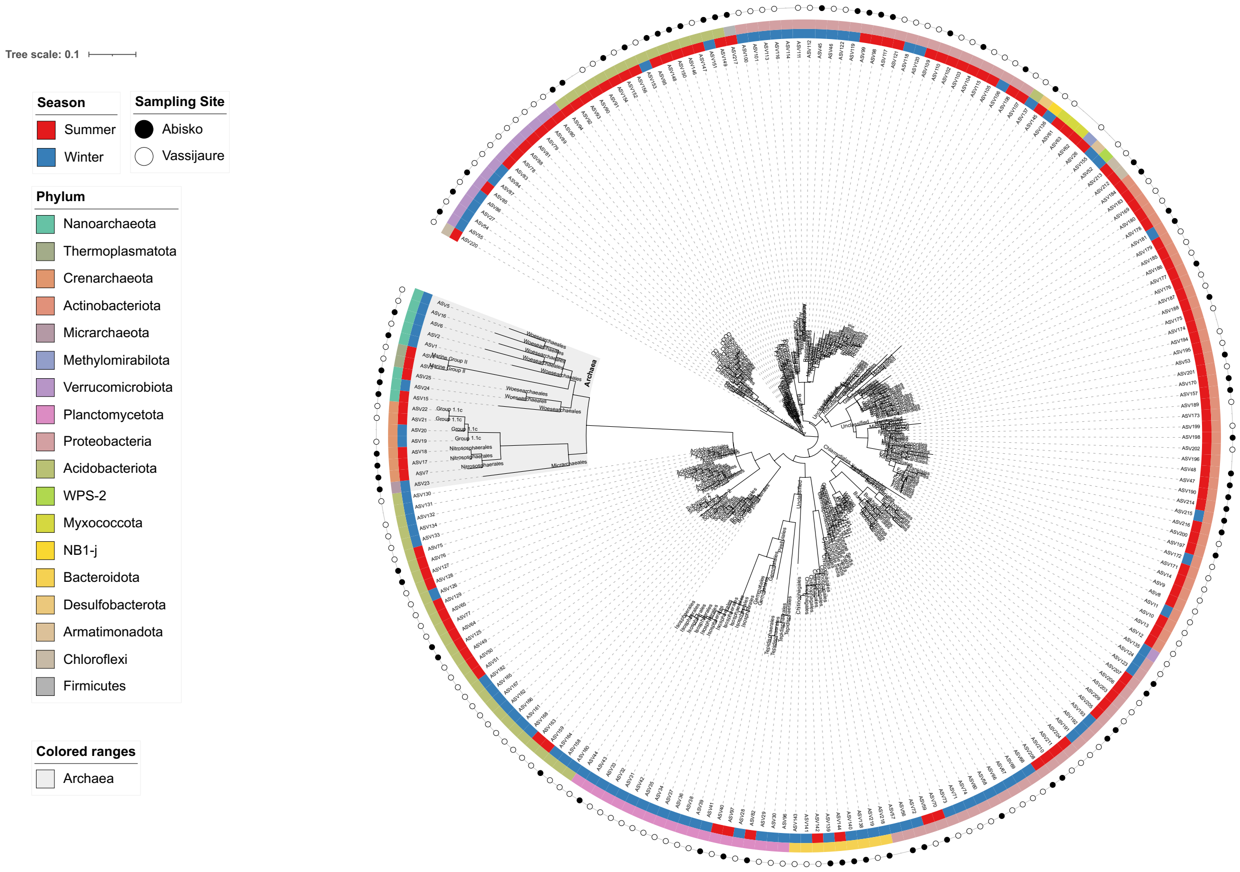


**Figure S6**: Approximate maximum-likelihood phylogenetic tree of seasonal bacterial and archaeal ASVs inferred using FastTree under a GTR model. The inner ring denotes ASV clustering as summer (red) or winter (blue) ASVs. Taxonomic classification on phylum level is denoted by the middle ring. Symbols in the outer ring denote sampling sites at which the ASVs were detected with missing symbols representing ASVs found at both sampling sites.

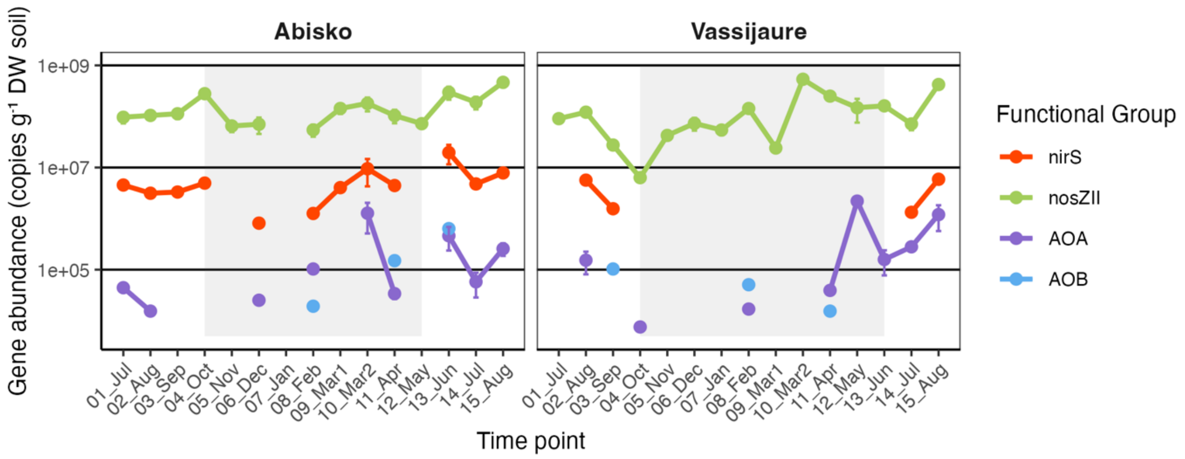


**Figure S7**: Variation in gene copy numbers for functional genes during one year from July 2019 to August 2020 at two tundra sites in Abisko and Vassijaure. Gene abundances are plotted on a $\log_{10}$ scale and show means with error bars denoting standard errors ($n \leq 5$). AOA – ammonia-oxidizing archaea (archaeal *amoA*), AOB – ammonia-oxidizing bacteria (bacterial *amoA*).
